# Distributed DNA-based Communication in Populations of Synthetic Protocells

**DOI:** 10.1101/511725

**Authors:** Alex Joesaar, Shuo Yang, Bas Bögels, Ardjan van der Linden, B.V.V.S. Pavan Kumar, Neil Dalchau, Andrew Phillips, Stephen Mann, Tom F. A. de Greef

## Abstract

Developing distributed communication platforms based on orthogonal molecular communication channels is a crucial step towards engineering artificial multicellular systems. Here, we present a general and scalable platform entitled ‘Biomolecular Implementation of Protocellular Communication’ (BIO-PC) to engineer distributed multichannel molecular communication between populations of non-lipid semipermeable microcapsules. Our method leverages the modularity and scalability of enzyme-free DNA strand-displacement circuits to develop protocellular consortia that can sense, process and respond to DNA-based messages. We engineer a rich variety of biochemical communication devices capable of cascaded amplification, bidirectional communication and distributed computational operations. Encapsulating DNA strand-displacement circuits further allows their use in concentrated serum where non-compartmentalized DNA circuits cannot operate. BIO-PC enables reliable execution of distributed DNA-based molecular programs in biologically relevant environments and opens new directions in DNA computing and minimal cell technology.

---

Living cells communicate by secreting diffusible signaling molecules that activate key molecular processes in neighboring cells^1,2^. These molecular communication channels facilitate information distribution among cells, enabling collective information processing functions that cannot be achieved by cells in isolation^3,4^. Synthetic biologists have advanced the engineering of synthetic cell-cell communication systems based on living cells resulting in multicellular consortia capable of complex sender-receiving functions^5-9^, bidirectional^10,11^ and synchronized^12^ communication and distributed computations^13,14^. However, engineering synthetic gene networks in living cells remains challenging due to the large number of context dependent effects arising from, among others, competition between shared resources and loading effects^15^. In contrast, developing molecular communication channels among abiotic synthetic protocell compartments has received much less attention than in living systems^16,17^. Due to their minimalistic design, engineering distributed information processing functions in fully synthetic multicellular communities has several advantages including a high degree of control and reduced design-build-test cycles. Abiotic protocellular consortia, based on lipid or non-lipid compartments and wired by orthogonal information channels, would thus present a versatile technology for the bottom-up construction of complex cell-population behaviors^18,19,20^. Although some elegant strategies to achieve one-way intercellular communication in fully synthetic protocellular systems have been reported^21,22,23,24^, a scalable methodology for implementing distributed functions involving bidirectional communication across populations of protocells is currently lacking.

Here, we present Biomolecular Implementation of Protocellular Communication (BIO-PC), a highly programmable protocellular messaging system that enables the construction of biochemical communication devices with collective information-processing functions. BIO-PC is based on protein-based microcapsules called proteinosomes^25^, containing internalized molecular circuits that encode and decode orthogonal chemical messages based on short single-stranded nucleic acids (Figure 1a). Initial experiments (Supplementary Fig. S1) revealed that proteinosomes, in contrast to liposomes, are permeable to short (<50 bp) ssDNA strands, making them highly suitable for the development of a protocellular communication platform. To code and decode ssDNA messages between individual protocells, we leveraged the modularity and scalability of dynamic DNA nanotechnology based on enzyme-free, toehold-mediated strand displacement.^26,27^ The high programmability and predictability of DNA strand-displacement (DSD) reactions allow the design of molecular circuits exhibiting a wide range of dynamic functions including catalytic cascades^28^, digital logic circuits^29^, Boolean neural networks^30^, distributed control algorithms^31^ and oscillations^32^; DNA is therefore well suited as a programmable molecular substrate. While molecular communication among localized enzyme-free^33,34^ and enzyme-driven^35^ DNA circuits has been reported, previous strategies were based on grafting DNA templates onto micrometer-sized beads. These examples elegantly show the possibilities of engineering collective behavior among amorphous agents; however, particle-based systems do not allow straightforward tuning of population dynamics and function suboptimally in biologically relevant media, since grafted DNA strands are directly exposed to the environment. Using BIO-PC, we experimentally demonstrate, for the first time, a scalable framework for distributed computation and bidirectional communication in populations of semipermeable microcapsules. We use microfluidic trapping devices to congregate protocellular communities capable of collective functions such as multiplex sensing, cascaded amplification, bidirectional communication and distributed logic operations (Figure 1b) and reveal how population dynamics can be easily tuned by controlling the compartment permeability. Finally, we show that encapsulating DNA gates inside proteinosomes makes them less vulnerable to digestion by nucleases, thereby greatly increasing their lifetime in concentrated serum and thus opening the way for the development of cell-like autonomous molecular sensors and controllers under physiological conditions.

**Figure 1:**
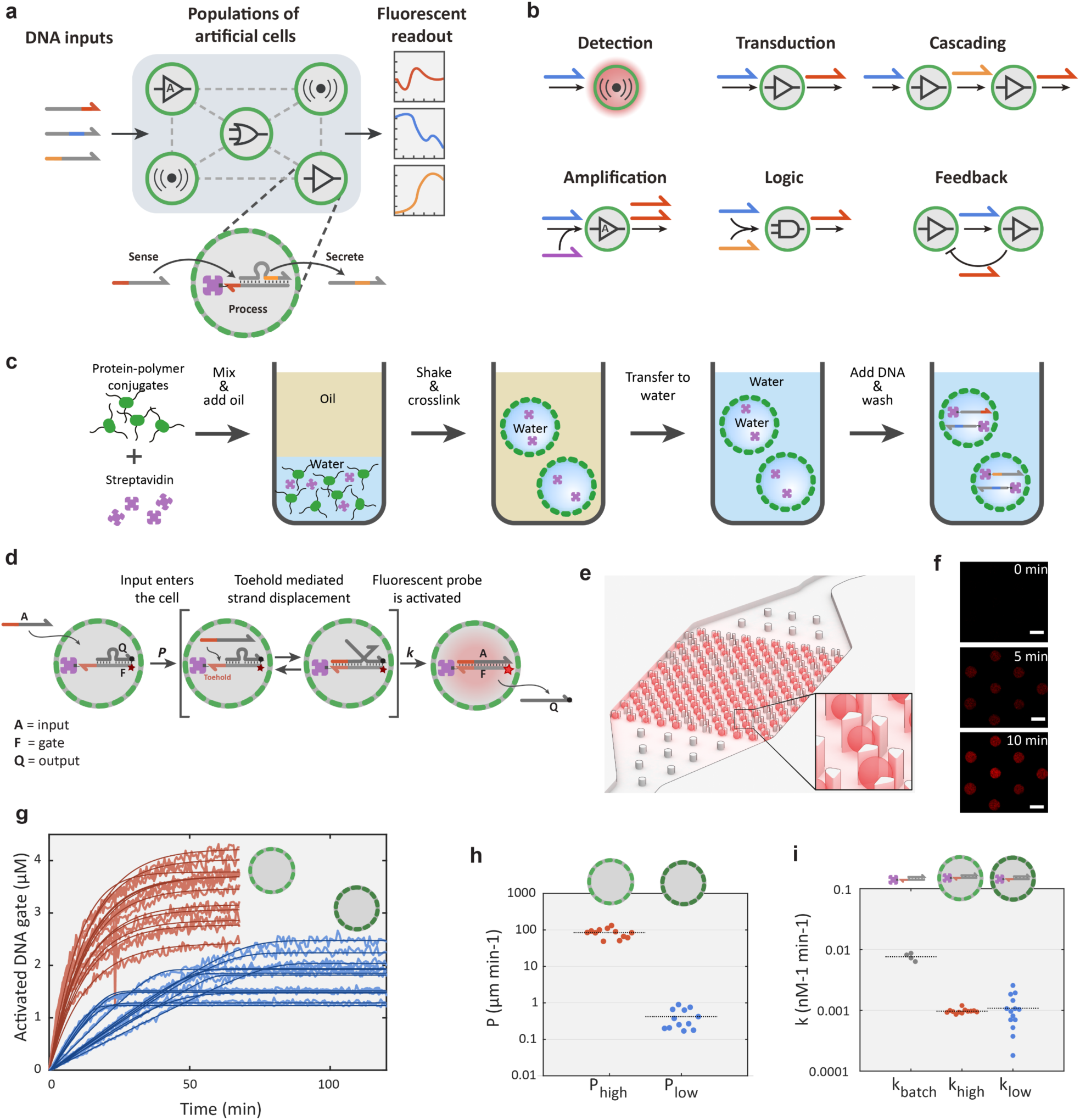
Design elements for biomolecular implementation of protocellular communication (BIO-PC). **a,** General strategy of the BIO-PC platform. Protocells with encapsulated DNA gate complexes are localized on a 2D spatial grid and can sense, process and secrete short ssDNA-based signals. The system is initiated by adding of ssDNA inputs and the response dynamics associated with the compartmentalized DSD reactions for each protocell are followed by confocal microscopy. **b**, Individual protocells can be configured to perform various tasks ranging from signal detection to Boolean logic operations. Individual modules can be combined to implement more complex population behaviors such as cascaded signaling, bidirectional communication and distributed computing. **c**, Procedure for preparing DNA-encapsulating protocells. Streptavidin-containing proteinosomes are assembled by covalently crosslinking BSA-NH_2_/PNIPAAm nanoconjugates at the interface of water-in-oil emulsion droplets. The crosslinked microcapsules are then phase transferred into water and biotinylated DNA strands are localized in the interior via biotin-streptavidin interactions. **d**, A mechanistic model for toehold-mediated DSD reactions inside protocells. The input strand **A** diffuses through the semipermeable membrane at a rate governed by permeability constant ***P*** (μm min^-1^) and then activates a fluorescent (Cy5) DNA gate complex **F:Q** (**F** = fluorophore/gate; **Q** = quencher/output strand) via a DSD reaction described by a bimolecular rate constant ***k*** (nM^-1^ min^-1^). **e**, CAD drawing of a microfluidic protocell trap array with computer-rendered trapped protocells shown in red. **f**, Confocal micrographs of eight trapped proteinosomes showing time-dependent increase in Cy5 fluorescence associated with the activation of an encapsulated DNA gate complex. Scale bar 50 μm. **g**, Fluorescence traces and model fittings of DSD reactions in protocells with high (red) and low (blue) membrane permeability. Reactions were carried out with 100 nM of **A** for high-***P*** and 1000 nM for low-***P*** proteinosomes. Concentrations of the activated DNA gate complex (**F:A** complex) are determined from time-dependent fluorescence measurements on individual protocells trapped within the microfluidic array device. **h**, Estimated permeability constants for the two protocell populations. **i**, Estimated bimolecular rate constants for the DSD reactions inside high-***P*** and low-***P*** proteinosomes compared to the estimated rate constant for a reference DSD reaction under batch conditions.

## Encapsulation and Remote Activation of DNA Circuits

To establish a highly programmable protocellular messaging system, we localized DNA gates inside protocells using the high-affinity, non-covalent interaction between biotin and streptavidin. Proteinosomes were prepared by mixing an aqueous solution of cationized bovine serum albumin (BSA-NH_2_)-poly(N-isopropylacrylamide) (PNIPAAm) nanoconjugates and streptavidin with oil, resulting in a Pickering emulsion where the amphiphilic BSA-PNIPAAm conjugates self-assemble at the interface, while the hydrophilic streptavidin preferentially remains in the aqueous interiors of the emulsion droplets (Fig. 1c). The self-assembled structures were chemically cross-linked using PEG-bis(N-succinimidyl succinate) (MW 2 kDa) and subsequently phase transferred into water via dialysis, resulting in polydisperse microcapsules (size range 10-60 μm) containing a distribution of encapsulated streptavidin (Supplementary Fig. S2 and S3). Confocal fluorescence imaging of FITC-labeled proteinosomes in combination with fluorescence recovery after photobleaching (FRAP) experiments revealed that the microcapsules have well-defined membranes consisting of cross-linked BSA polymer amphiphiles and liquid-like interiors that contain excess BSA conjugate and streptavidin (Supplementary Fig. S4). Earlier studies have determined that proteinosomes have porous membranes with a molecular weight cutoff *ca.* 40 kDa^25^. Using fluorescently labeled streptavidin (MW 52.8 kDa), we verified that the encapsulation of streptavidin inside the proteinosomes is stable for more than a month (Supplementary Fig. S3). We can expect that in addition to the size, which is slightly above the cutoff, streptavidin is also weakly crosslinked to some extent by the crosslinker, further assisting its stable encapsulation for such long periods of time. Proteinosomes containing encapsulated DNA were prepared by adding short (MW < 20 kDa), biotinylated ssDNA strands that can diffuse through the semipermeable membrane and anchor to the encapsulated streptavidin (Supplementary Fig. S5). Excess DNA was removed by sedimentation and washing, yielding proteinosomes containing internalized DNA with an overall loading efficiency between 30-50% (Supplementary Fig. S5) and which can be stored at 4 °C for more than a month without significant DNA loss (Supplementary Fig. S5).

To assess the feasibility of implementing DSD-circuits inside proteinosomes, we started by characterizing the performance of a simple toehold-mediated strand-displacement reaction^36^. An inactivated DNA gate complex (**F:Q**), consisting of a fluorophore (Cy5)-modified gate strand (**F**) with an exposed toehold domain and a quencher-labeled output strand (**Q**) was localized inside the proteinosomes by binding to encapsulated streptavidin (Fig. 1d). To quantitatively analyze reaction kinetics, we designed a microfluidic platform capable of single-protocell capture, activation and imaging (Fig. 1e and Supplementary Fig. S6). In addition to providing control over protocell population density and restricting their movement, the device removes abnormally large and small proteinosomes, allowing analysis of a more homogeneous population of protocells (Supplementary Fig. S2). The protocells were loaded into the trap array by flowing a proteinosome suspension through the trapping chamber until a desired number of protocells had been captured (Methods). To initiate DSD within the captured protocells, a membrane-permeable complementary input strand (**A**) was flowed into the microfluidic device, resulting in an increase in Cy5 fluorescence for each protocell over time due to displacement of **Q** and formation of the activated **F:A** complex (Figure 1f and 1g, red lines). Importantly, a control experiment employing a fluorescently-labeled input strand with a non-complementary toehold domain showed that DNA strands can diffuse across the proteinosome membrane but fail to displace the output strand to a considerable level (Supplementary Fig. S5). Together, these results establish that encapsulated DNA gates are activated via a two-step process, in which input strands first diffuse across the protocell membrane and subsequently initiate DSD reactions.

Compared to other types of localized DNA circuits on surfaces^37^, microparticles^35^ or DNA origami^38,39^, encapsulating DNA circuits inside proteinosomes introduces membrane permeability constant (***P***) as an additional parameter in the reaction kinetics. We therefore explored whether we could alter membrane permeability to tune the overall dynamics of the DSD reaction. Using a shorter crosslinking agent (BS(PEG)5) significantly reduced the rate of the proteinosome-encapsulated DSD reactions (Fig. 1g, light blue traces). Comparing the experimental traces shows that DNA gate activation dynamics are markedly different between the high-and low-***P*** proteinosomes. While high-***P*** proteinosomes display exponential DNA gate activation kinetics, typical _f_or non-compartmentalized DSD reactions^36^, low-***P*** proteinosomes show linear kinetic profiles until high levels of DNA gate activation are attained. To rationalize the influence of membrane permeability on the strand-displacement kinetics, the input strand **A** was labeled with Alexa546. This allowed us to simultaneously observe ditffusion of **A** into the protocells as well as intra-protocellular activation of the **F:Q** DNA gate by toehold mediated displacement of the quencher strand (Supplementary movie 1, Supplementary Fig. 5). We first estimated the value of ***P*** for each individual proteinosome assuming Fickian diffusion (Fig. 1h). The analysis shows that the permeability constant for the input strand is approximately two orders of magnitude smaller in the low-***P*** proteinosomes. Using the estimated permeability constants and a bimolecular approximation for the DSD reaction^36^, we constructed a simple ODE model that describes the time-dependent activation of the encapsulated DNA gate – considering input diffusion from the inter-protocellular environment (Methods, Supplementary Fig. 7) – and obtained estimates for the bimolecular rate constant ***k*** for each protocell. The simple two-parameter kinetic model describes the experimental traces very well for both proteinosome types (Fig. 1f, dark red and dark blue traces) and, importantly, reveals that both populations have similar strand-displacement rate constants. These results indicate that the rate-determining step in low-***P*** proteinosomes is the input diffusion across the membrane, while in high-***P*** proteinosomes it is the DSD reaction. While the estimated DSD rate constants are similar for both the high-and low-***P*** proteinosomes, they are almost an order of magnitude lower compared to a DSD reaction under batch conditions (Fig. 1i). The lower DSD rate constant of the encapsulated DNA gates can be attributed to a difference in local chemical environment (higher local DNA gate concentrations, presence of soluble nanoconjugates, for example) compared to the same reaction under batch conditions. Collectively, these findings establish BIO-PC as a versatile platform that allows compartmentalized DNA gates to be actuated by complementary DNA-based messages, the dynamics of which can be tuned by protocellular membrane permeability.

## Cascaded amplification of DNA messengers among protocell communities

Having established the feasibility of performing compartmentalized DSD reactions inside proteinosome-based protocells, we proceeded to determine whether the BIO-PC platform could be used to implement chemical communication among different proteinosome populations. Intercellular amplification of soluble messengers among different populations of cells is a characteristic feature of complex multicellular systems such as the immune system. To this end, we designed an inter-protocellular non-enzymatic DNA-based signaling cascade based on two proteinosome populations containing different streptavidin-anchored DNA gate complexes (Fig. 2a and Supplementary Fig. S8). The two proteinosome populations were loaded into the trap array sequentially, resulting in a random spatial arrangement of the binary population (Methods). An externally added input strand diffuses into the protocells resulting in toehold-mediated displacement of a quencher-labeled output strand specifically in protocell population **1** which can be observed from the concomitant increase in Cy5 fluorescence associated with formation of an activated gate complex in population **1**. Secretion of the released output strand (**Q_1_**) by population **1** functions as a chemical signal that is sensed by population **2** (Supplementary Fig. S8). Detecting the signal strand via toehold-mediated strand displacement of a quencher labeled output strand (**Q_2_**) leads to higher Alexa546 fluorescence due to activation of the DNA gate complex in the second population of protocells.

**Figure 2:**
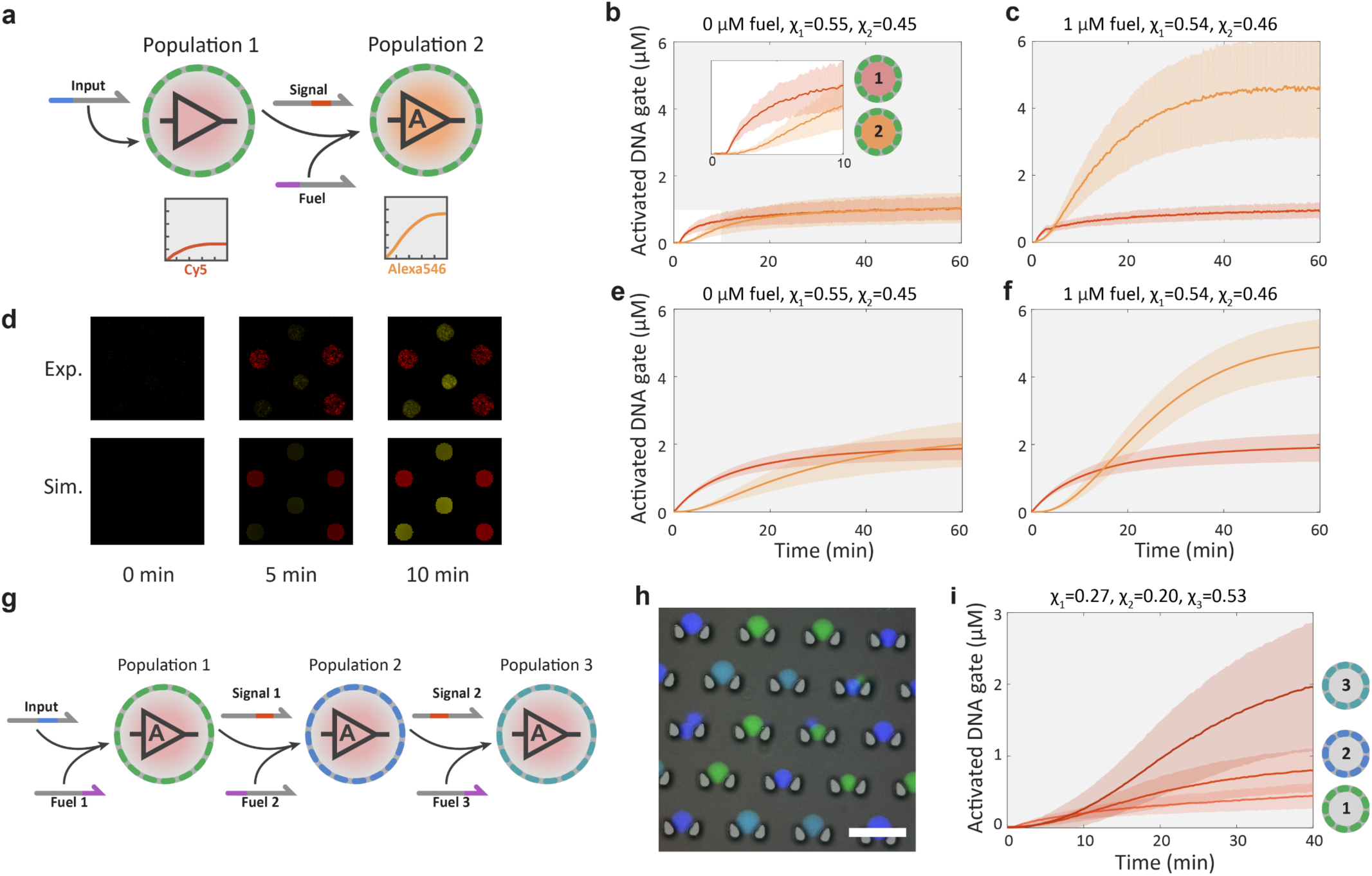
Signaling cascades between protocell populations using non-enzymatic DNA signal amplification. **a,** Abstract diagram of a two-layer signaling cascade between two protocell populations. Proteinosomes of the first population sense the externally added input strand and respond by activating a Cy5 fluorescent DNA gate complex and secreting a signal strand to the external environment. The second population can sense the signal and produces a fluorescent response. Non-enzymatic DNA catalysis is used to recycle the signal strand by consuming the abundant fuel strand, thereby producing an amplified response. **b**, Mean and standard deviations the fluorescent traces of two high-***P*** proteinosome populations in the absence of a fuel strand. [Input] = 100 nM, the fractions of the two populations were χ_1_=0.55, χ_1_=0.45. Concentrations of the activated DNA gate complexes are determined from time-dependent Cy5 (population **1,** red profile) or Alexa546 (population **2,** orange profile) fluorescence measurements on individual protocells trapped within the microfluidic array device. **c**, Mean and standard deviations the fluorescent traces of two high-***P*** proteinosome populations in the presence of 1 μM of fuel strand. [Input] = 100 nM, χ_1_=0.54, χ_1_=0.46. Note the large increase in the fluorescent output of population **2** (orange profile) due to signal amplification**. d**, Confocal micrographs (Exp.) and simulation data (Sim.) of a group of six proteinosomes comprising three members of population **1** and three members from population **2.** The delay between activation of population **1** and **2** during the initial minutes of the reaction is shown. **e,** Mean and standard deviations of 2D simulation data using parameters of the system in **b** (red and orange curves, population **1** and **2**, respectively). **f**, Simulation data using parameters of the system in **c. g,** Abstract diagram of three-layer signaling cascade with each stage providing fuel-mediated signal amplification upon activation by its cognate input. **h**, Epifluorescence micrograph of three-color barcoded high-***P*** proteinosomes overlaid on a bright-field micrograph of the microfluidic trapping array. The barcoding is realized by membrane labelling of proteinosome populations **1**, **2** and **3** using FITC (green), DyLight405 (dark blue) and a 1:1 mixture of the two fluorophores (light blue), respectively. **i**, Cy5 fluorescence traces of proteinosome populations **1**, **2** and **3** of the three-layer signaling cascade showing progressive levels of signal amplification. Reaction was performed using 10 nM of input and 500 nM of each fuel strand.

While protocells of population **1** implement a simple transducer functionality, where one input strand activates a single fluorescent DNA gate complex and leads to the secretion of one signal molecule, protocells of the second population can amplify their input signal. We implemented signal amplification by employing non-enzymatic DNA catalysis, which recycles the signal strand after its initial strand-displacement reaction using abundant fuel strands^29,40^. Without signal amplification (Fig. 2b), the system response depends on the relative fraction of the two populations, with a 1:1 ratio resulting in approximately equal levels of DNA gate complex activation. When 1 μM fuel strand is added to the system (Fig. 2c), population **2** can produce a much stronger response, and the final value depends on the concentration of encapsulated DNA gate complex determined by the streptavidin loading (Supplementary Fig. S8). Importantly, as is expected from a cascaded network, a delay between the responses of the two populations is clearly visible (Fig. 2b inset, Fig. 2d and Supplementary Movie 2). The response characteristics of the binary population can also be tuned by changing the protocell membrane permeability, with low-***P*** proteinosomes of population **1** providing a gradual linear response and significantly delaying the activation of population **2** (Supplementary Fig. S8). To confirm the experimental observations, we simulated the two-population system using the Visual DSD^41^ software by solving the corresponding reaction-diffusion equations in 2D (Methods). The numerical model considers the full complexity of DSD reactions in each of the two populations and DNA signal diffusion between identical-sized protocells on a grid (Fig. 2d). To simulate the observed variability in the dynamic behavior of the protocells, the initial DNA gate complex concentration inside each compartment was drawn from a random distribution that matched the experimental distribution. The model was parametrized using the average DSD rate constant and permeability obtained from the analysis of the single-population DSD experiments. As can be observed from Figures 2e and f, the mathematical model can capture the most important features of the experimental data including the steady-state values, the time delay between the two populations and the effect of the fuel. While the experimental variability in dynamic behavior between protocells is partly captured by the model, additional effects such as the size distribution of protocells and variance in protocell permeability are expected to further increase inter-protocell variability.

To further demonstrate the scalability and orthogonality of the BIO-PC platform, we daisy-chained three protocell populations, each capable of strand transduction and fuel-induced signal amplification upon activation by its cognate input (Fig. 2g and Supplementary Fig. S9). To circumvent the need for three spectrally non-overlapping fluorescent dyes to monitor strand-displacement, we used a barcoding method where the protocell membrane was labeled with FITC, DyLight405 or a 1:1 mixture of the two (Fig. 2h), allowing a single fluorophore (Cy5) to monitor all three DSD reactions. Upon addition of 10 nM of input strand and in the presence of 500 nM of each fuel strand, each population can produce an amplified response with respect to the previous population (Fig. 2i). Control experiments (Supplementary Fig. S9) in which either population **2** or input strand is not present reveal that population **3** at the end of the chain only displays a low autoactivation caused by signal leakage inherent to catalytic DSD reactions^42^. Together, these results establish BIO-PC as a versatile platform to program diffusively connected compartmentalized cascades and reveal the high orthogonality of DNA-based molecular messengers.

## Bidirectional Communication and Distributed Sensing and Computing in Protocell Populations

In living systems, intercellular communication via diffusible factors is often bidirectional and includes both positive and negative regulatory interactions^43^. In addition, distributed computation across multiple cells is central to complex multicellular systems ranging from the immune system^44^ to neural networks. Here we use the BIO-PC platform to show the implementation of bidirectional communication and distributed sensing and processing of DNA-based inputs. The synthetic version of a bidirectional feedback loop (Fig. 3a and Supplementary Fig. S10) is based on two high-***P*** proteinosome populations where the activation of population **1** on addition of an input strand results in the secretion of a signal strand and a concomitant increase in Cy5 fluorescence. The secreted signal can activate protocells of population **2** which then respond with an increase in Alexa546 fluorescence and secretion of an orthogonal inhibitor strand that can deactivate population **1** by displacing the input strand and re-quenching the Cy5 fluorophore. Protocells of population **2** also implement an amplification module, allowing them to recycle the input strands. The results demonstrate that after the input strand is added, population **1** is rapidly activated followed by a more gradual activation of population **2** and the simultaneous deactivation of population **1** (Fig. 3b and Supplementary Movie 3). Decreasing the level of the input and fuel strands dampens the response in population **1** and delays its peak activation level; however, the pulse length remains unchanged (Fig. 3c). In the absence of population **2**, protocells in population **1** remain active (Supplementary Fig. S10), confirming that negative feedback signaling occurs in the binary population. Without the input strand, the amplification module of population **2** exhibits low levels of leakage, while population **1** shows no response. (Supplementary Fig. S10).

**Figure 3:**
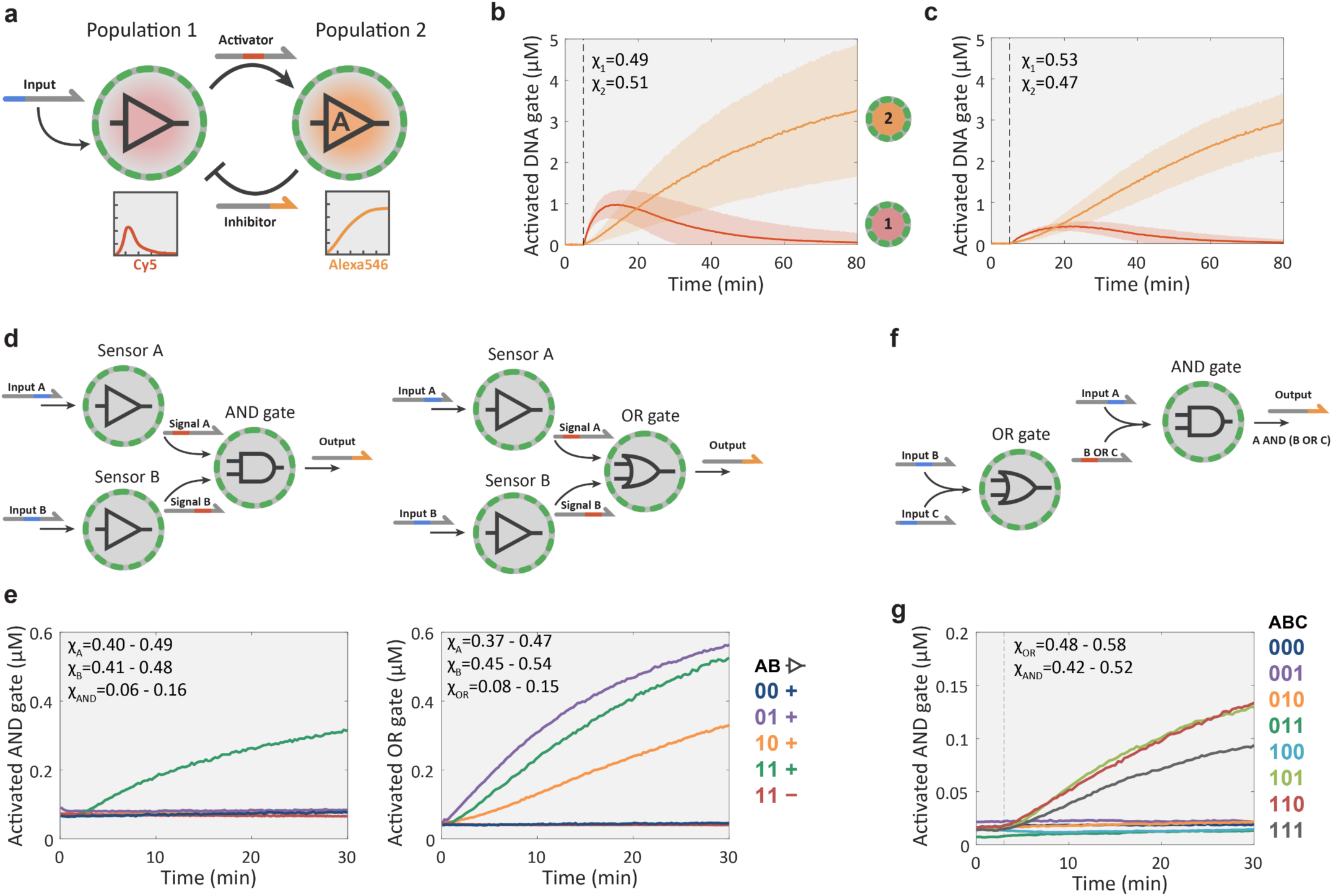
Feedback signaling and Boolean logic. **a,** Abstract diagram of a negative feedback loop between two protocell populations engaged in compartmentalized DSD reactions. The system is triggered by introducing the ssDNA input strand into the external medium, which then activates the Cy5-labeled DNA gate complex of population **1** and induces the release of a strand that acts as an activator for population **2**, switching on their Alexa546-labeled DNA gate complex. As a consequence, an inhibitor strand is released into the medium and protocells in population **1** are deactivated by switching off their fluorescent DNA gate complexes. **b**, Time-dependent activation levels for two high-***P*** proteinosome populations implementing the negative feedback loop. The activation and deactivation of population **1** (red curve, Cy5 fluorescence) is consistent with negative feedback signaling in the binary population. The progressive increase in the fluorescence of population **2** (orange curve, Alexa546) is due to fuel-mediated signal amplification. The reaction was initiated with 200 nM of input and 1 μM of fuel strands to run the amplification reaction in population **2**. The fractions of the two populations were χ_1_=0.49, χ_2_=0.51. **c**, Activation levels with reduced levels of input (100 nM) and fuel (300 nM), χ_1_=0.53, χ_1_=0.47. **d**, Schematic representations of combinatorial sensing-and-processing protocell network with either AND or OR logic gates. See Supplementary Figure S11. **e**, Experimental data of the AND and OR devices based on a protocell network consisting of two sensor proteinosome populations in combination with an AND or OR gated population. The plots show time-dependent changes in the concentration of the activated DNA gate complex in the AND or OR proteinosomes. The color-coded traces correspond to the configurations indicated in the truth table, with the first two columns indicating the presence (1) or absence (0) of the input strands A and B respectively and third column indicating the presence (+) or absence (-) of the sensor populations. All populations comprise high-***P*** proteinosomes with equal input strand concentrations (100 nM) where applicable; In experiments with all three populations, their respective fractions were within the range of χ_A_=0.40-0.49, χ_B_=0.41-0.48 χ_AND_=0.06-0.16 for the AND gate and χ_A_=0.37-0.47, χ_B_=0.45-0.54 χ_OR_=0.08-0.15 for the OR gate. The complete data with standard deviation for each input configuration and the activation of the individual sensor modules are shown on Supplementary Fig. S11. **f**, Schematic representation of two high-***P*** proteinosome populations collectively implementing a serially gated logic circuit. Population **1** calculates logic OR from inputs **B** and **C**, and the protocells of population **2** calculate logic AND of input **A** and the output of population **1**. See Supplementary Figure S12. **g**, Experimental data of the two-gate, three-input logic circuit. The color-coded traces correspond to the input configurations shown in the truth table with 1 indicating the presence and 0 the absence of the respective input strand. Experiments were performed with 500 nM of each input strand, χ_OR_=0.48-0.58, χ_AND_=0.42-0.52. The complete data sets with standard deviation for each input are shown in Supplementary Fig. S12.

The orthogonality of DSD circuitry makes it an excellent platform for implementing Boolean logic operations^28,29^. When combined with spatial segregation of components, it should be possible to design and construct amorphous computing devices capable of combinatorial sensing and distributed signal processing. As a proof of principle, we used the BIO-PC platform to implement a combinatorial sensing-and-processing device based on a community of three protocell populations comprising a network of two sensor populations with simple transducer functionality and one logic-gate population with either 2-input AND or OR function (Fig. 3d, Supplementary Fig. S11). Detailed kinetic characterization of the two multi-protocellular consortia revealed that the sensor modules are activated by their cognate input (Supplementary Fig. S11) and reported OR and AND computations according to the truth table (Fig. 3e). To implement more complex computational functions, we designed a distributed computing device consisting of a serially layered OR and AND logic-gate population and showed that the system responds to the correct input patterns (Fig. 3f and g, Supplementary Fig. S12).

Population based DSD reactions are inherently less controlled than batch-operated DSD experiments, with run-to-run variability introduced by variation in the total number of protocells, ratios between the populations and their spatial distribution, and we therefore observe run-to-run variation in the collective outputs for the TRUE input configurations. However, for both sensing-and-processing devices and the distributed computing device we observe that, despite variation in conditions and output levels, the TRUE and FALSE states are always clearly distinguishable. While these sensing and processing functions can also be implemented using non-compartmentalized DSD circuits, orthogonal strand-level design is critical and typically requires several optimization cycles to lower unwanted cross-hybridization and leakage reactions. The modular nature of BIO-PC, however, permits the use of non-orthogonal strands as DNA gates are physically separated. Indeed, under batch conditions the two modules of the serial logic circuit cross-hybridize and thereby render it inoperative (Supplementary Fig. S12). Collectively, these results reveal that feedback control and complex computational operations can be engineered by a library of simple protocell modules communicating via orthogonal DNA messengers.

## Operation of DNA Circuits in a Biologically Relevant Environment

One of the long-standing goals of DNA nanotechnology is to create autonomous molecular machines that can operate in harsh biological environments and provide drug-delivery or diagnostic functionality^45^. Recent studies have shown that the functionality of simple DSD circuits is severely limited in cell culture medium with 10% fetal bovine serum (FBS) ^46^. We performed a series of experiments to investigate whether proteinosomes could be used to protect DSD networks from degradation in a biological environment. First, simple DSD circuit stability was tested under batch conditions (Fig. 4a). A streptavidin-anchored DNA gate complex (**F:Q**) at a total concentration of 100 nM was incubated in 50% FBS for 48h, after which a cognate strand (**A**) was added and the fluorescence associated with formation of the activated **F:A** complex was measured. The results show that the FBS-incubated DNA gate complex lost nearly all of its activity whilst the gate complex remains fully operational when incubated for the same amount of time in an aqueous buffer (Fig. 4b). Initial experiments revealed that proteinosomes prepared using our existing method would abruptly result in osmotic collapse when placed into 50% FBS (Supplementary Fig. S13). We therefore proceeded to increase the mechanical rigidity of proteinosome by adding 60 mg/mL of nonfunctionalized BSA to the interior of the protocells (Methods). The resulting proteinosomes were stable in FBS and had more gel-like interiors, as evidenced by FRAP analysis (Supplementary Fig. 13). When the DNA gate complexes were now encapsulated inside proteinosomes (Fig. 4c) and subjected to the same 48h incubation in 50% FBS, the system retained nearly all of its activity compared to incubation in 0% FBS (Fig. 4d). Even though the local DNA gate concentration inside proteinosomes is higher than the DNA gate concentration used in the batch experiments, the ratio of total DNA to FBS was far higher in the latter (Supplementary Methods) confirming the protective effect associated with compartmentalization of the DSD reactions within the proteinosomes. Additionally, we tested whether localizing the DNA gate complexes to polystyrene beads similar in size to the proteinosomes would facilitate a protective effect and found that while some activity was retained after incubation in 50% FBS for 48h, most likely due to the steric hindrance of the nucleases (Supplementary Figure S13). The activity associated with the beads was considerably lower than a control experiment where the incubation was undertaken in a buffer solution (Supplementary Figure S13).

**Figure 4:**
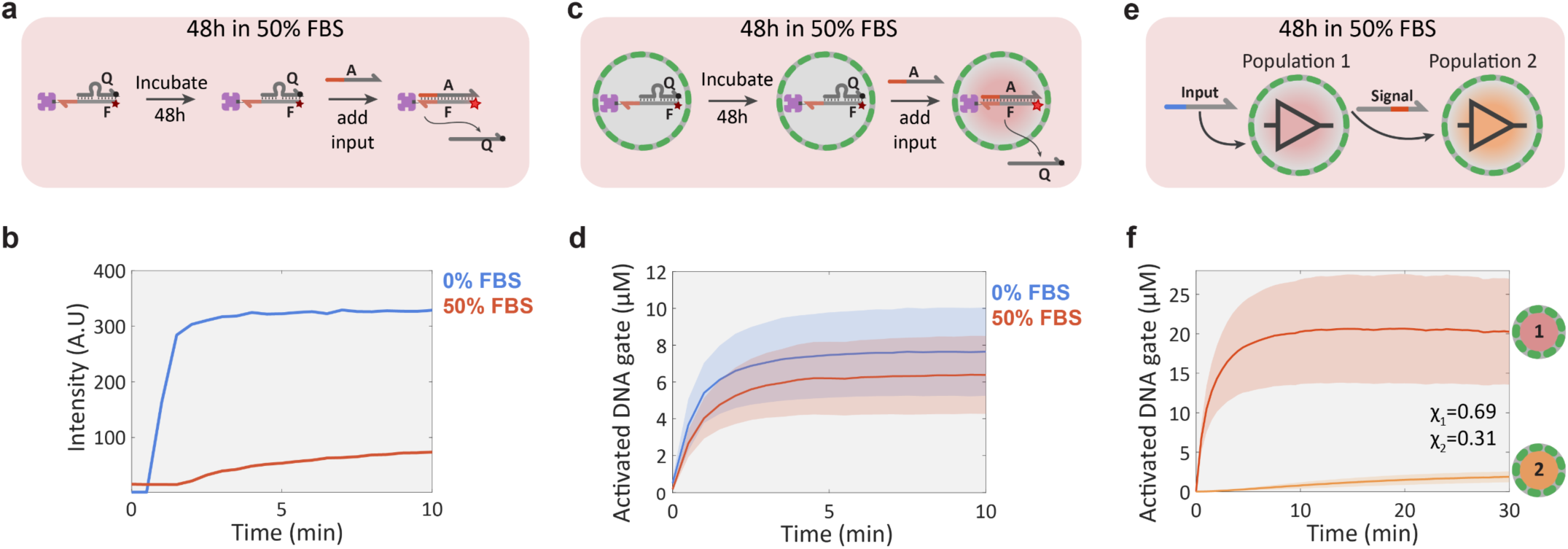
Compartmentalized DNA reaction networks in 50% FBS. **a**, Experimental procedure for validating the stability of a streptavidin-bound quenched DNA gate complex (**F:Q**, 100 nM; streptavidin, 100 nM) in 50% FBS under batch conditions. The stability of the complex was determined by measuring the fluorescence associated with DSD activity after addition of input strand **A** (200 nM). **b**, Fluorescence traces for the batch DSD reaction in 0% (buffer only, blue trace) and 50% (red) FBS showing loss of activity in 50% FBS. **c**, Validation procedure for a compartmentalized version of the DSD reaction shown in **a**. **d**, Mean and standard deviation of protocell activation levels in 0% (buffer, blue trace) and 50% (red) FBS. The experiments were performed by incubating the DNA gate-containing proteinosomes for 48h in 50% FBS or 0% FBS, followed by adding 2 μM of input strand **A** to initiate the DSD reaction. High-***P*** proteinosomes containing free BSA (60 mg/mL) were used to minimize osmotic collapse. **e**, Schematic representation of a simple non-catalytic transducer-receiver signaling cascade between two protocell populations incubated for 48 h in 50% FBS. Proteinosomes of population **1** sense the externally added input strand and respond by activating a Cy5 fluorescent DNA gate complex and secreting a signal strand that is sensed by population **2** to produce a Alexa546 fluorescent response. **f**, Mean and standard deviations of protocell activation levels for populations **1** and **2** of the 2-stage signaling cascade showing fluorescent activation in both populations. The reaction was started after 48h incubation in 50% FBS by adding 2 μM of the input strand. High-***P*** proteinosomes containing free BSA (60 mg/mL) were used to minimize osmotic collapse.

To determine the feasibility of DNA-based inter-protocellular communication in a biologically relevant environment, we constructed a simple signaling cascade between two populations (Fig. 4e). The binary protocell population was incubated in 50% FBS for 48h, placed into a microfluidic trap array and then activated by addition of 2 μM of input strand. All steps were performed in 50% FBS without any intermittent washing steps using aqueous buffers. The results show an immediate fluorescence response in the transducer population **1** associated with displacement of the quencher strand, followed by a slow increase in the fluorescence read-out from population **2** (Fig. 4f). Approximately 2 μM of the DNA gate in the receiver population **2** is activated within 30 minutes (Fig. 4f). Although the activity in population **2** is significantly reduced (Supplementary Fig. 13), most likely due to signal strand degradation during transit through the serum-containing external environment, the results show that DNA signal transmission between the two protocell populations is possible in concentrated serum. These results confirm that encapsulation of DSD circuits in proteinosomes enables their operation in a biologically relevant environment and demonstrate the possibility of distributed DNA-based communication under challenging conditions.

## Conclusions

Biological systems employ diffusion-based molecular communication channels to distribute tasks and collectively adapt to their environment. Here we developed BIO-PC, a modular protocellular communication platform based on non-living components and demonstrate the successful design and characterization of a range of distributed collective behaviors. Our strategy is based on the streptavidin-mediated encapsulation of DNA templates inside protein-based semipermeable microcapsules that encode and decode molecular messages using DNA-based strand displacement reactions. Because the information processing components are physically separated from each other, well-characterized and high-performing parts can be reused. This important feature could, for example, aid the design of multi-layer DNA-based winner-take-all circuits^47^ capable of classifying complex information patterns. Such circuits would be highly valuable in disease diagnostics^48^.

BIO-PC allows a high degree of control over molecular communication by providing a tunable barrier, a feature that makes it distinct from previous methods based on tethering rule-encoding DNA templates to surfaces. Gines *et al*. have recently shown^35^ that in the absence of a physical boundary, the emergence of collective behavior in bead-based communities is severely constrained as a result of competition among local chemical reactions and global diffusive transport as described by the dimensionless Damkohler number. While we have not performed a full mathematical analysis on the competition between local and global effects, our experimental results reveal that it is possible to independently tune the diffusive transport rate by preparing proteinosomes using different chemical cross-linkers (Figure 1g, Supplementary Figure S7 and S8). Proteinosome porosity could also be addressed using selectively degradable building blocks^49^ or changing the chemical composition of the protein-polymer nanoconjugate. Furthermore, proteinosomes also present unique possibilities to engineer feedback between molecular communication and protocell permeability which is not possible when using bead-based systems. For example, a recently published clamped hybridization chain reaction (C-HCR)^50^ in which a single stranded DNA-based initiator induces the 3D self-assembly of two metastable hairpins could be used to selectively promote gelation within the interior of proteinosomes upon receiving a DNA-based instruction.

While protocell-scale systems do not display the same levels of information-processing properties as living cells, their minimal and streamlined design could yield simple multicellular model systems with a high degree of control and the potential to uncover generalizable concepts^18,19^ In order to simplify the analysis of such protocell-protocell circuit motifs, it is important that the variability in the dynamic response of proteinosome-based protocells can be controlled. This could be achieved by preparing proteinosome micro-droplets by microfluidic methods that has been shown to result in proteinosome populations with narrower size distributions^51^ than those observed when using standard bulk methodologies, as employed in this study. Living cells utilize a high level of temporal control and a wide range of non-linear regulatory strategies, such as competition, shuttling and degradation of soluble factors, to tune intercellular communication dynamics^4,43^. The versatility of DNA-based chemical reaction networks allows implementation of these regulatory mechanisms by employing DNA-based threshold gates^29^, time-delay circuits^52^ or exonuclease-assisted degradation of DNA messengers^35^. Adding these functions to BIO-PC would facilitate the construction of a large library of protocell-protocell circuit motifs and a systematic analysis of their temporal behavior. We anticipate that scalable architectures for distributed computation across populations of communicating protocells could enable complex information processing tasks for future diagnostic and therapeutic applications.

## Acknowledgements

We thank Georg Seelig, Willem Mulder, Leroy Cronin and Pascal Pieters for helpful discussions. This work was supported by the European Research Council, ERC (project n. 677313 BioCircuit), ERC Advanced Grant Scheme (EC-2016-ADG 740235) and Marie Curie Individual Fellowship, an NWO-VIDI grant from the Netherlands Organization for Scientific Research (NWO, 723.016.003), funding from the Ministry of Education, Culture and Science (Gravity programs, 024.001.035 & 024.003.013) and BrisSynBio (BB/L01386X/1).

## Author contributions

A.J. designed the study, performed experiments, analyzed the data and wrote the manuscript. S.Y. and B.B. performed experiments and analyzed the data. A.L. and A.J. designed and fabricated the microfluidic chip. N.D. and A.P. performed computational experiments. P.K. and S.M. provided key reagents and provided critical input for the initial experiments. T.G. conceived, designed and supervised the study, analyzed the data and wrote the manuscript. S.M. revised the manuscript. All authors discussed the results and commented on the manuscript.

## Methods

### Synthesis BSA-NH_2_/PNIPAAm nanoconjugates

Cationized BSA (BSA-NH_2_) was synthesized according to a previously reported method^25^. Typically, a solution of diaminohexane (1.5g, 12,9 mmol in 10 mL of MilliQ water) was adjusted to pH 6.5 using 5 M HCl and added dropwise to a stirred solution of BSA (200 mg, 3 μmol in 10 mL of MilliQ water). The coupling reaction was initiated by adding 100 mg of 1-(3-Dimethylaminopropyl)-3-ethylcarbodiimide HCl (EDC) immediately and another 50 mg after 5h. If needed, the pH value was readjusted to 6.5 and the solution was stirred for a further 6h and then centrifuged to remove any precipitate. The supernatant was then dialyzed (Medicell dialysis tubing, MWCO 12-14 kDa) overnight against MilliQ water and freeze-dried.

End-capped mercaptothiazoline-activated PNIPAAM (M_n_ = 9800 g mol^-1^, 4 mg in 5 mL of MilliQ water) was synthesized according to the previously reported method^24^ and added to a stirred solution of BSA-NH_2_ (10 mg in 5 mL of PBS buffer at pH 8.0). The solution was stirred for 10 h and then purified using a centrifugal filter (Millipore, Amicon Ultra, MWCO 50 kDA), and freeze-dried.

FITC-and DyLight405-labeled BSA-NH_2_/PNIPAAm conjugates were prepared using the same method, except that labeled BSA (Supplementary Methods) was used as the starting material.

### Preparing streptavidin-containing proteinosomes

In a typical experiment, BSA-NH_2_/PNIPAAm conjugates (final concentration: 8 mg mL^-1^), streptavidin (final concentration: 1, 4, or 10 μM) and 1.5 mg of PEG-bis(N-succinimidyl succinate) (Mw = 2000, approximately 38 PEG units on average) (or alternatively 2 mg of BS(PEG)5 to prepare proteinosomes with a lower permeability for ssDNA strands (Low-***P*** proteinosomes)) were mixed in 30 μL of 50 mM sodium carbonate buffer (pH 9). 500 μL of 2-ethyl-1-hexanol was immediately added and the mixture was shaken by hand for 20 s to produce a Pickering emulsion. After 3 h of sedimentation, the upper clear oil layer was discarded, 800 μL of 70% ethanol was added and the emulsion was gently shaken. The dispersion was then sequentially dialyzed against 70% and 50% ethanol for 2 h each and finally against Milli-Q water for 24 h. The proteinosomes were then stored at 4°C for later use.

To produce fluorescently labeled proteinosomes, a 1:7 mixture of labeled (FITC, DyLight 405 or a 50/50 mixture of the two)/unlabeled BSA-NH_2_/PNIPAAm conjugates was used. The proteinosomes used in the FBS experiments were prepared using the same method, except that non-functionalized BSA (final concentration 60 mg/mL) was added to the aqueous phase and 2 mg of PEG-bis(N-succinimidyl succinate) was used as the crosslinking agent.

### DNA sequence design

DNA strands were designed by hand with the help of a MATLAB script that generates random sequences with a desired fraction of each nucleotide. The design of the reactive DNA gates localized inside proteinosomes was mainly based on the “seesaw” gate motif of Qian and Winfree^29^. Instead of using separate gate:output and reporter complexes, the fluorescent reporting functionality was built into the gate:output complex itself by labeling the gate strands (denoted as **F**) with a fluorophore (Cy5, Cy3 or Alexa546) and the output strands (denoted as **Q**) with a quencher. All the gate strands have two chemical modifications – a biotin-TEG to facilitate binding to encapsulated streptavidin, and a fluorophore to monitor the DSD reaction. The two modifications were positioned at the opposite ends of the gate strands. In the initial inactive state, the gate strands were hybridized to complementary output strands, which had a quencher modification so that the fluorescence of a gate:output complex was quenched. We used a complementary input strand (**A**) to activate the gate by displacing the output strand via a toehold-mediated strand-displacement reaction, resulting in an active gate:input complex where the fluorophore is unquenched. In some experiments, the output strand of one gate was used as an input for another gate, thereby functioning as a signal strand. In addition, networks were designed to implement signal amplification using a fuel strand that reacted with the active gate:input complex to displace the input strand, yielding an active gate:fuel complex. The released input strand reacted with another gate:output complex, thereby functioning as a catalyst. The molecular reaction diagrams for all the DSD networks can be found in Supplementary Figures S8 to S11. The sequences and chemical modifications of all strands can be found in Supplementary Tables 1-6. DNA strands for the 3-stage cascade network, negative feedback and logic gates were designed by following a set of rules based on earlier studies^28^: A three-letter nucleotide code was used for all strands (A, T, and G were used for the gate strands and A, T, and C for input, output, and fuel strands). No more than four consecutive A or T nucleotides (except for nonfunctional poly-T domains at the biotin end of certain gate strands that function as spacers), and no more than three consecutive C or G nucleotides were employed. All strands had a G (or C) fraction between 0.3 and 0.7. All sequences were tested with NUPACK^53^ to detect any possible undesired interactions and redesigned if necessary.

### DNA oligonucleotide synthesis

All DNA oligonucleotides were purchased from Integrated DNA Technologies with HPLC purification. 100 μM and 10 μM stock solutions were made using nuclease free TE buffer (10 mM Tris, 0.1 mM EDTA, pH 8.0, Integrated DNA Technologies) and stored at −20°C.

### Localization of DNA gate complexes in streptavidin-containing proteinosomes

The buffer for all experiments, unless otherwise specified, was 10 mM Tris (pH 8.0, Invitrogen) with 12 mM Mg^2+^ (Invitrogen) and 0.1 % v/v Tween 20 (Sigma). In a typical localization experiment 10 μL of a dispersion of streptavidin-containing proteinosomes, 5 μL of 4X buffer and 2 μL of biotinylated DNA gate strand (from a 10 μM stock solution) were gently mixed with a pipette in a 1.5 mL Eppendorf tube and incubated at room temperature for 1 h, followed by addition of 3 μL output strand (from a 10 μM stock solution), gentle mixing and overnight incubation at 4°C. The excess unbound output strand was removed as follows: 10 μL of the supernatant was carefully removed from the top and discarded, 400 μL of buffer was added and the proteinosomes were resuspended by mixing with a pipette. The proteinosomes were allowed to sediment for 1 to 2 h or alternatively spun down using a microcentrifuge (4k RCF, for 5 min) and 400 μL of supernatant was removed from the top and discarded. This process was repeated and the resulting suspension of streptavin/DNA gate complex-containing proteinosomes stored at 4°C.

### Design and fabrication of microfluidic trapping devices

We designed a two-layer microfluidic chip to facilitate the physical trapping and *in situ* imaging of populations of streptavidin/DNA-loaded proteinosomes. The chip (Supplementary Fig. S6) consisted of a 1.5 mm X 2 mm or 1.5 mm X 1.5 mm localization chamber with PDMS pillars, a filtering chamber, inlet channels with pneumatically actuated Quake style push-up valves^54^ and an outlet channel. Master molds for the two layers were fabricated on separate silicon wafers (Silicon Materials) using standard photolithography techniques^54^. The molds for bottom and top layers were made by spin-coating SU8-3050 to a height of 50 μm, and spin-coating AZ 40xt to a height of 40 μm, respectively. After development the AZ 40xt mold was reflowed, resulting in rounded channels with a height of ∼60 μm at the center. The microfluidic chips were assembled from PDMS using standard multilayer soft lithography techniques^54^ and plasma bonded to circular #1.5 glass coverslips.

### Proteinosome trapping and activation

The microfluidic trapping device was mounted on the stage of a confocal laser scanning microscope (CLSM, Leica SP8). Control channels were filled with MilliQ water and actuated using a pneumatic valve array (FESTO), which was in turn actuated using a programmable logic controller (PLC, WAGO Kontakttechnik GmbH). The PLC was connected to the Ethernet port of a PC and controlled using a custom Matlab GUI. The pressure in the control channels was 2 bar. The pressure to the inlet channels was controlled using adjustable pressure regulators (Flow-EZ, Fluigent). In a typical experiment, buffer solution was connected to inlet port 1. First, air bubbles were pushed out of the flow channels by pressurizing the buffer channel at 1 bar and closing all other inlet and outlet valves, followed by thoroughly washing all the flow channels using the buffer solution. Next streptavidin/DNA-containing proteinosomes were loaded into the trap array from inlet port 2 at a pressure of 10 mbar. The inlet port 2 was then closed and the proteinosomes were gently washed (the exact pressure needed to achieve enough flow for effective washing, but without forcing the protocells out of the traps depended on the fluidic resistance of the microfluidic setup and was experimentally determined beforehand; 5 to 15 mbar was normally used) with the buffer solution for 5 to 10 min to remove any unbound DNA. For multi-population experiments, in which it was critical that the unbound DNA was reduced to the minimum to avoid leakage reactions, the populations were loaded sequentially with intermittent washing periods. To ensure that protocells of different populations were well mixed, additional mixing steps were performed when necessary. Mixing was achieved by first flowing the prococell suspension in reverse direction into one of the input channels and then back into the trapping chamber, this process randomizes the spatial distribution of protocells from different populations. The confocal microscope was focused on the trapping chamber and time-lapse imaging was started. All the experiments were performed at room temperature. The initial steady-state signals were recorded as the baseline values.

The input (and fuel, if used) DNA oligonucleotides were diluted (and mixed) to the desired concentration in buffer solution with 2 mg/mL BSA (Molecular Biology Grade, New England Biolabs) to minimize DNA adsorption to tubing and pipette tips and loaded to inlet port 3. The DSD reactions were started by flowing the input DNA into the trapping chamber using 10 mbar pressure.

### Incubation and activation of proteinosomes in concentrated fetal bovine serum (FBS)

Proteinosomes (5 μL) with localized DNA gate complexes and loaded with non-functionalized BSA (60mg/mL added during assembly), 4X buffer solution (5 μL) and FBS (10 μL) were mixed in a 1.5 mL Eppendorf tube and incubated at room temperature for 48h. The localization of the proteinosomes in a microfluidic trap array was performed as described in the previous section, except that FBS was added (50% final v/v) to the washing buffer and also to the input DNA solution.

### Data acquisition and analysis

Fluorescence data were acquired using a confocal laser scanning microscope (CLSM, Leica SP8) equipped with solid state lasers (405 nm for DyLight405, 488 nm for FITC, 552 nm for Cy3 and Alexa546, 638 nm for Cy5) and a hybrid detector. The time-lapse measurements were performed with either a 10X/0.40NA (1.55X1.55mm field of view, 7 μm slice thickness), or a 20X/0.75NA objective (0.775X0.775mm field of view, 2 μm slice thickness) at a resolution of 512X512 pixels. The photon counting mode of the hybrid detector was used. Concentration values of the activated DNA gate complexes were obtained from the relative fluorescence units (RFU) measured by the CLSM using the following procedure: First, the RFU values were corrected by subtracting the baseline value (baseline correction was not performed for the data of the logic gate circuits on Fig. 3e and g to give a more objective representation of the experimental variation between the different input configurations). The baseline-corrected values were then converted to activated DNA gate complex concentrations using calibration values of the corresponding activated gate complex. In order to rule out any N-quenching^55^ effects due to the input or fuel strand binding to the gate strand, all calibrations were performed using the corresponding gate:input or gate:fuel complexes instead of using free gate strands. Calibrations were performed by filling the microfluidic trapping chamber with 1 μM of the respective activated DNA gate complex, capturing a fluorescent image and recording the average RFU value over a straight line across the device (Supplementary Fig. S14). The linearity of the detector and the method itself were verified by measuring the fluorescence of the activated DNA gate complex over a range of know concentrations (Supplementary Fig. S14).

### Mathematical model and parameter estimation of proteinosome permeability and compartmentalized DSD reactions

Using Fick’s first law, the diffusion of a DNA oligonucleotide *A* into a proteinosome can be described by the following ODE (its derivation is given in Supplementary Note 1):

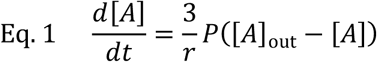

where [*A*] is the strand *A* concentration inside the protocell, [*A*]_out_ is the strand *A* concentration outside the protocell, ***P*** (μm min^-1^) is the permeability constant of the proteinosome membrane and *r* is the radius of the proteinosome. The permeability constants in Fig. 1h were obtained by observing a fluorescently labeled strand *A* diffuse into a proteinosome population and measuring the concentrations ([*A*]_out_ and [*A*]) over time. For the low-***P*** proteinosomes the permeability constant was the rate-limiting step and could be determined from an experiment with DSD (Fig. 1d) by assuming all *A* that had entered the protocell reacted with the gate complex on a timescale much faster than the diffusion process. Thus, the gradient of *A* across the membrane can be assumed to be constant (equal to [*A*]_out_) at the initial linear phase of the reaction and the permeability constant can be calculated by solving 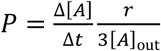 (Supplementary Figure S7). For the high-***P*** proteinosomes, the diffusion process occurred at a much faster rate. Because it was not possible to separate the diffusion process from the DSD reaction as they occurred at a similar timescale, a different method was used to estimate the permeability constant. After the DSD reaction (Fig. 1d) had completed, the remaining free *A* was washed away, followed by an additional injection of *A*. The diffusion of *A* was measured over time inside and outside the proteinosomes, and the permeability constant was estimated by non-linear least squares optimization of equation 1 using nonlinear least-squares solver (lsqnonlin, MATLAB) (Supplementary Figure S7).

To estimate the DSD rate constants a model (shown graphically on Fig. 1d) consisting of the following ODEs was used.

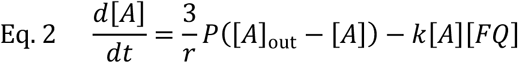

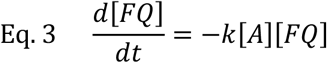

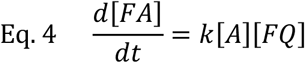

where [*FQ*] is the concentration of the inactive gate complex, [*FA*] is the concentration of the active gate complex and *k* (nM^-1^ min^-1^) is the forward rate constant of the strand displacement reaction.

The rate constants *k*_high_ and *k*_low_ in Fig. 1i were obtained by measuring the activated DNA gate complex concentration ([*FA*]) over time and performing parameter estimation using a nonlinear least-squares solver (lsqnonlin, MATLAB) with the previously estimated ***P*** values as constants.

### 2D reaction-diffusion simulations

Computational simulations of the 2-stage signaling cascade (Fig. 2d, e and f) were performed using the Chemical Reaction Network (CRN) tool of Visual DSD^41,56^, a software tool for designing and analyzing DNA strand displacement networks. The Command Line Interface (CLI) of the tool was used and the configuration file was generated using a custom MATLAB script. A 2D model was used for the simulations; as in the experimental setup, the proteinosomes were placed on a 2D grid and the fluorescence data obtained from a 2D slice. Proteinosome positions were assigned based on the CAD drawing of the microfluidic device (Supplementary Fig. S6), and the population (1 or 2) was randomly assigned for every proteinosome with a 1:1 ratio between the two populations. The encapsulated gate complex concentrations were sampled from a normal distribution based on the experimental measurements of the mean and variance of the encapsulated DNA gate concentrations of the respective proteinosome populations. The radius, permeability and DSD rate constant were set to be the same across all the proteinosomes in order to simplify the mathematical model. The diffusion coefficient of the DNA oligonucleotides in the aqueous buffer was estimated using an empirical formula *D*_*W*_ = 4,9 * 10^-6^cm^2^/s ∗ [bp size]^−0.72^ from an earlier study^57^. The complete description of the mathematical model and computer code is given in Supplementary Note 2. System simulation was performed by solving the reaction-diffusion equations with the CRN tool.

## Data availability

The data that support the findings of this study are available from the corresponding author, by a reasonable request.

