## Supporting Information for "Distributed DNA-based Communication in Populations of Synthetic Protocells"

[1.] Laboratory of Chemical Biology, Department of Biomedical Engineering, Eindhoven University of Technology, The Netherlands.
[2.] Institute of Complex Molecular Systems, Department of Biomedical Engineering, Eindhoven University of Technology, The Netherlands.
[3.] Computational Biology Group, Department of Biomedical Engineering, Eindhoven University of Technology, The Netherlands.
[4.] Centre for Protolife Research and Centre for Organized Matter Chemistry, School of Chemistry, University of Bristol, Bristol BS8 1TS, UK.
[5.] Microsoft Research, Cambridge CB1 2FB, UK.

#### Supporting Information: Methods, Movies, Figures, Notes, Tables

### Supplementary Methods

#### Materials

2-Ethyl-1-hexanol (Sigma, 98%), 1-(3-dimethylaminopropyl)-3-ethylcarbodiimide HCl (EDC, Carbosynth), 1,6-diaminohexane (Sigma, 98%), Fluorescein isothiocyanate (FITC, 90%, Sigma), PEG-bis(N-succinimidyl succinate) (Mw = 2000, Sigma), streptavidin from *Streptomyces avidinii* (Sigma), DyLight™ 405 NHS ester (ThermoFisher), BS(PEG)5 (Sigma), streptavidin, Alexa Fluor™ 546 conjugate (ThermoFisher), bovine serum albumin (heat shock fraction, pH 7, ≥98%, Sigma), AZ 40xt (MicroChemicals), SU8 3050 (MicroChemicals), polydimethylsiloxane (PDMS, Sylgard 184), fetal bovine serum (Sigma) were used as received.

#### Labeling of BSA with fluorescent dyes

BSA was labeled with fluorescein isothiocyanate (FITC) as follows: 200 mg of BSA was dissolved in 10 mL of 50 mM sodium carbonate buffer (pH 9). 2.36 mg of FITC was dissolved in 590 µL of DMSO and added to the stirred BSA solution. The solution was stirred for 5 h, purified by dialyzing (Medicell dialysis tubing, MWCO 12−14 kDa) overnight against MilliQ water and freeze-dried.

BSA was labeled with DyLight 405 as follows: 30 mg of BSA was dissolved in 6 mL of 50 mM sodium carbonate buffer (pH 9). 1 mg of DyLight 405 NHS ester was dissolved in 100 µL of DMF and added to the stirred BSA solution. The solution was stirred for 2 h, purified by dialyzing (Medicell dialysis tubing, MWCO 12−14 kDa) overnight against MilliQ water and freeze-dried.

#### Fluorescence measurements on a plate reader

Batch testing of the serial logic circuit (Supplementary Fig. S12) was performed on a Tecan Spark 10M plate reader using 384 well plates (Nunc) with a reaction volume of 80 µL. Excitation and emission wavelengths were 635 nm and 680 nm respectively. Fluorescence measurements were performed every 20 s with an integration time of 40 µs.

#### DNA strand displacement reaction on streptavidin-functionalized polystyrene beads in FBS

Streptavidin-functionalized polystyrene beads in the size range of 18.0-25.0 µm functionalized with approximately $3.09\times10^{7}$ streptavidin molecules per bead were used (Spherotech). In a typical experiment 2 µL of bead dispersion, 5 µL of 4X buffer, 8 µL water and 2 µL of biotinylated DNA gate (**F**) strand (from a 0.122 µM stock solution) were gently mixed with a pipette in a 1.5 mL Eppendorf tube and incubated at room temperature for 1 h, followed by addition of 3 µL quencher/output (**Q**) strand (from a 0.122 µM stock solution), gentle mixing and overnight incubation at 4°C. This DNA/bead ratio yields beads with approximately $8.4*10^{-11}$ µmol of DNA localized per bead, which is comparable to the total amount of DNA in a similarly sized proteinosome with an inner DNA concentration of 20 µM. The DNA sequences are given in Supplementary Table 6. The removal of the unbound output strand was performed as follows: 10 µL of the supernatant was carefully removed from the top and discarded, 400 µL of buffer was added and the beads were resuspended by mixing with a pipette. The beads were allowed to sediment for 1 to 2 h or alternatively spun down using a microcentrifuge (4k RCF, for 5 min) and 400 µL of supernatant was removed from the top and discarded. This process was repeated for one more time and the resulting beads with localized DNA gates were stored at 4°C.

The DNA-functionalized beads were incubated in an Eppendorf tube with 50 % FBS for 48 h. The activation reaction was performed by mixing 2 µL of the suspended beads with 2 µL of 2 µM input DNA (**A**). The mixture was placed on a covered glass slide and images were obtained using a confocal laser scanning microscope (CLSM). As the beads were functionalized only on the surface (Supplementary Figure 13), the RFU level for each bead was obtained by averaging the RFU level over a circle around the periphery of the bead where the fluorescence level was the highest. A control experiment was performed where the incubation took place in a buffer solution in place of 50 % FBS.

#### Supplementary Movies

**Supplementary Movie 1 | Diffusion of a fluorescently labeled input DNA strand into protocells and activation of a fluorescent DNA gate complex.** Time lapse confocal microscopy recording with the green, red and yellow channels showing the fluorescence of FITC-labeled BSA-PNIPAAm building blocks, Cy5-labeled localized fluorescent DNA gate complex (**F:Q**), and Alexa546 labeled input ssDNA (**A)** respectively. This video corresponds to the reaction shown in Fig. 1d and g (blue traces – low-***P*** proteinosomes) in the main text and Supplementary Figure 7a-c. At t = 2 min, input strand (A) is flowed into the microfluidic trapping chamber and can be seen to gradually diffuse into and accumulate within the proteinosomes, where it displaces the quencher/output strand (**Q**) from the inactive DNA gate complex (**F:Q**), resulting in the activated DNA gate complex (**F:A**) and thereby in increased Cy5 fluorescence.

**Supplementary Movie 2 | Activation of the two-layer signaling cascade.** Time lapse confocal microscopy recording with the left channel showing the FITC fluorescence and the right channel showing the combined fluorescence of the Cy5 labeled DNA gate complex ($\mathbf{F_1:Q_1}$) of population **1** and the Alexa546 labeled DNA gate complex ($\mathbf{F_2:Q_2}$) of population **2**, of the network shown on Fig. 2a and Supplementary Fig. 8a. The experiment was performed using high-***P*** proteinosomes (prepared using 4 µM and 10 µM of streptavidin in population **1** and **2** respectively), with 1 µM of fuel strand and 100 nM of input strand.

**Supplementary Movie 3 | Negative feedback loop in two populations.** Time lapse confocal microscopy recording with the left channel showing the FITC fluorescence and the right channel showing the combined fluorescence of the Cy5 labeled DNA gate complex ($\mathbf{F_1:Q_1}$) of population **1** and the Alexa546 labeled DNA gate complex ($\mathbf{F_2:Inh}$) of population **2**, of the network shown on Fig. 3a and Supplementary Fig. 10. The experiment was performed using high-***P*** proteinosomes (prepared using 4 µM and 10 µM of streptavidin in population **1** and **2** respectively), with 1 µM of fuel strand and 200 nM of input strand.

#### Supplementary Figures

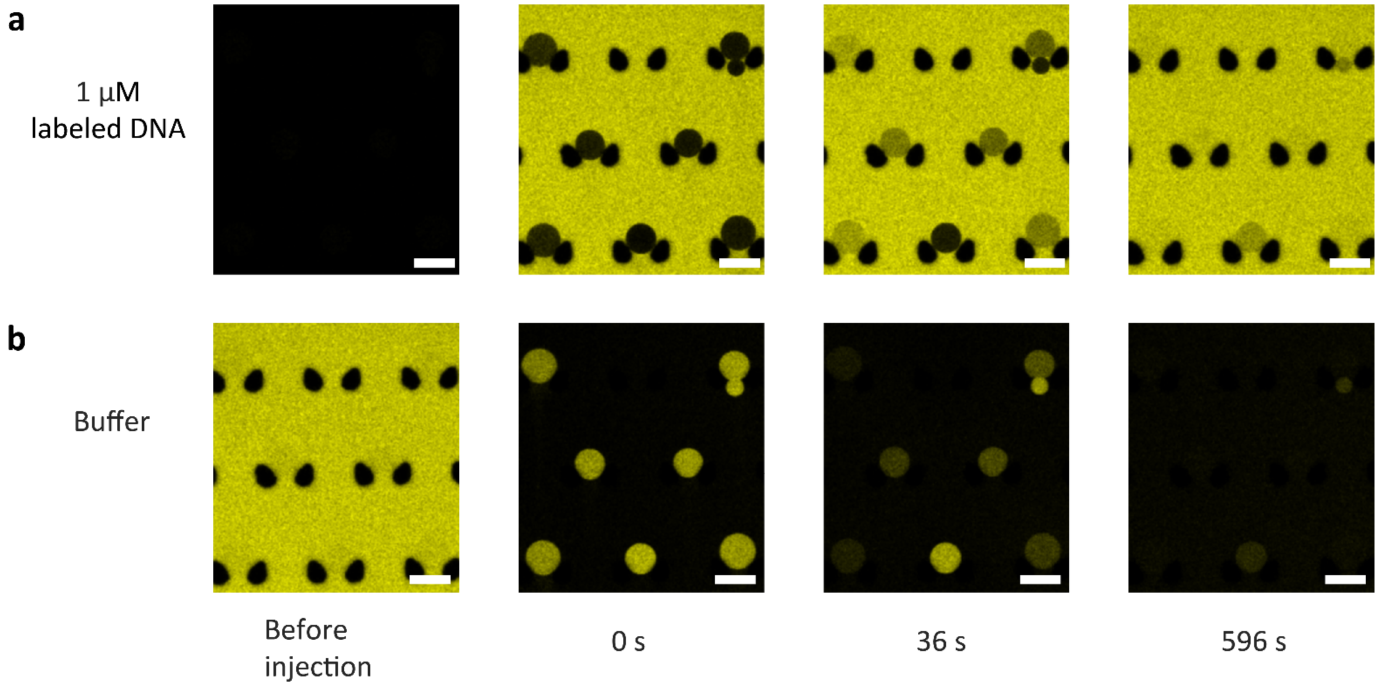


**Supplementary Fig. 1 | Diffusion of fluorescently labeled DNA oligonucleotides through proteinosome membrane. a**, Time lapse confocal micrographs of fluorescently labeled ssDNA (A_Alexa546, Supplementary Table 1) diffusing into the interior of individual proteinosomes captured in a microfluidic trapping device. The ssDNA does not have a biotin modification and the only driving force for its diffusion into the protocells is the concentration gradient across the membrane. **b**, Time-lapse confocal micrographs showing diffusion of fluorescent *ss*DNA out of proteinosomes. Samples were incubated as in **a** for 1 hour to equilibrate the internal and external concentrations, followed by removing *ss*DNA in the exterior by washing with a buffer solution and monitoring the decrease in fluorescence associated with the proteinosome interior Scale bars are 50 μm.

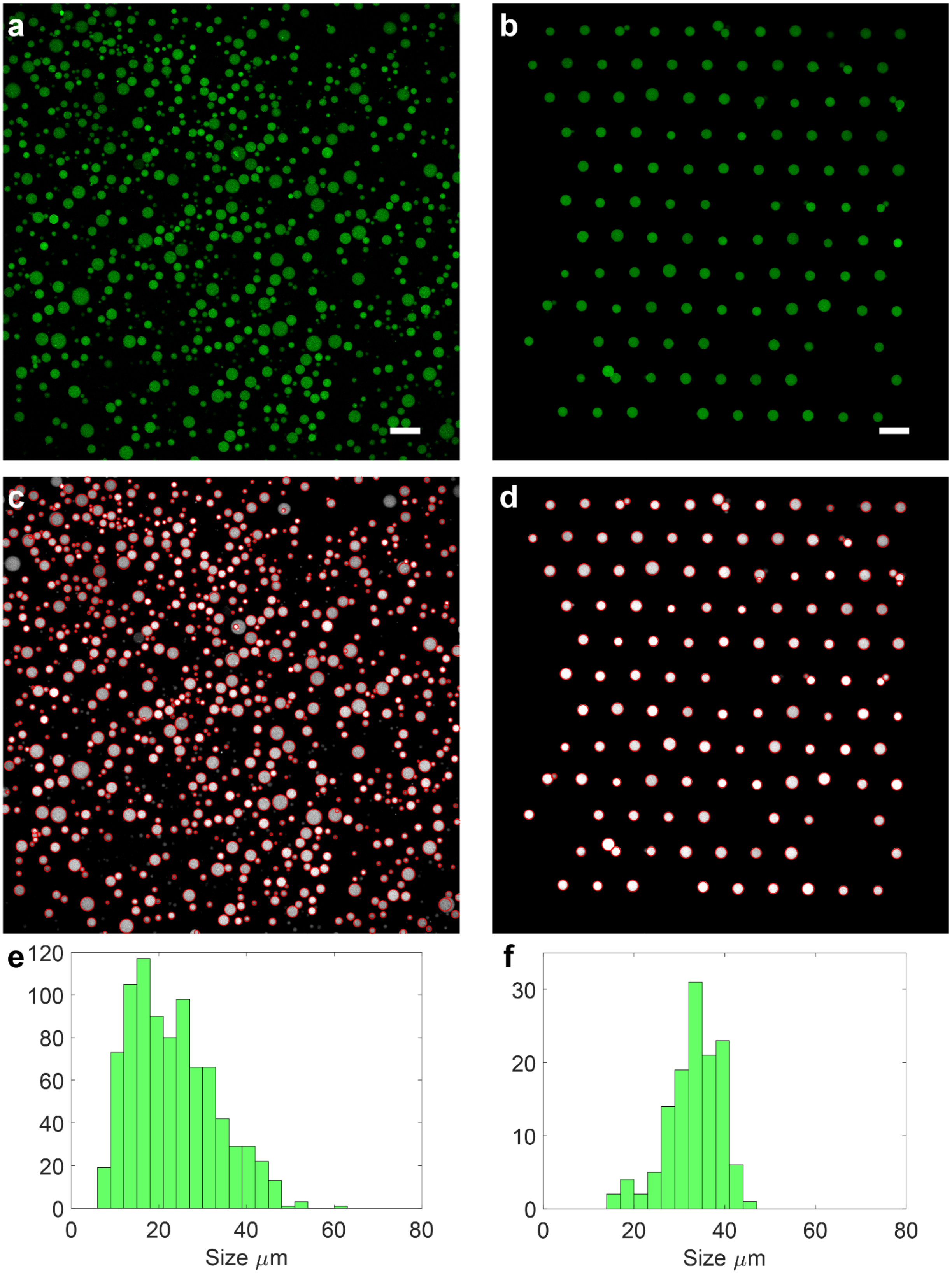


**Supplementary Fig. 2 | Comparison of proteinosome size distributions. a**, Confocal micrograph of FITC labeled high-**P** proteinosomes in an aqueous solution on a glass slide. **b**, Proteinosomes from the same batch as in **a** localized in a microfluidic trapping device. Scale bars are 100 µm. **c** and **d**, Processed images with red circles indicating the detected proteinosomes used for size distribution calculation. **e** and **f**, Histograms of the diameters of the detected proteinosomes.

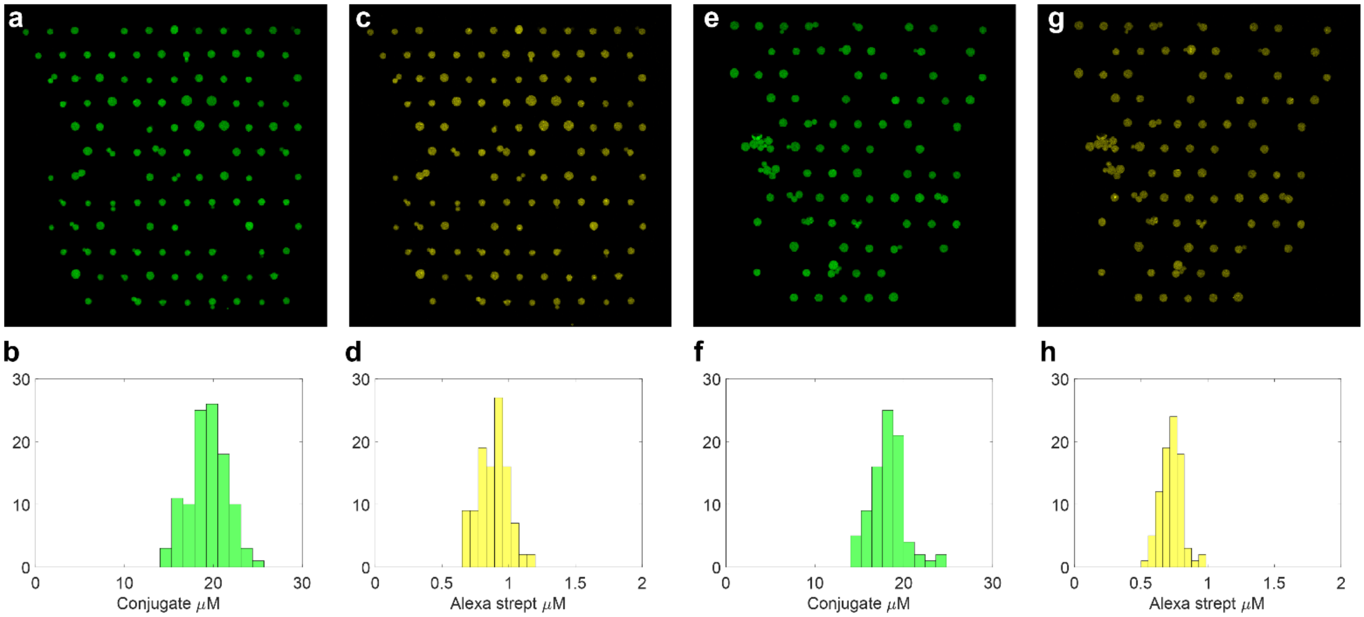


**Supplementary Fig. 3 | Distributions of proteinosome-encapsulated FITC labeled BSA-$NH_2$/PNIPAAm conjugate and Alexa546 labeled streptavidin. a,** Confocal micrograph of the FITC channel of trapped FITC labeled high-**P** proteinosomes with encapsulated Alexa546 labeled streptavidin. . The proteinosomes were prepared using excess BSA-$NH_2$/PNIPAAm (8 mg/mL, approx. 100 µM) such that the water-filled interior contained *ca.* 20 µM of the free conjugate; this increased the rigidity and ease of manipulation of the proteinosomes in the microfluidic trapping device. FITC was excited using 488 nm laser and the detection window was 498 nm – 528 nm. **b**, Histogram showing the distribution of the conjugate concentration in the interior of the protocells shown in **a**. **c**, Confocal micrograph of the Alexa546 channel showing the labeled encapsulated streptavidin. Alexa546 was excited using 552 nm laser and the detection window was 562 nm – 602 nm. **d**, Histogram showing the distribution of the streptavidin concentration in the interior of the protocells shown in **c**. 4 µM of streptavidin was used in the aqueous phase during the proteinosome assembly, resulting in an encapsulation efficiency of around 25%. **e – h**, The same measurements as in **a-d** performed one month after the proteinosomes were assembled and stored at 4° C in water. On an average > 80% of the encapsulated streptavidin and > 90% of the encapsulated conjugate had been retained. These values do not account for the potential degradation of the fluorophores.

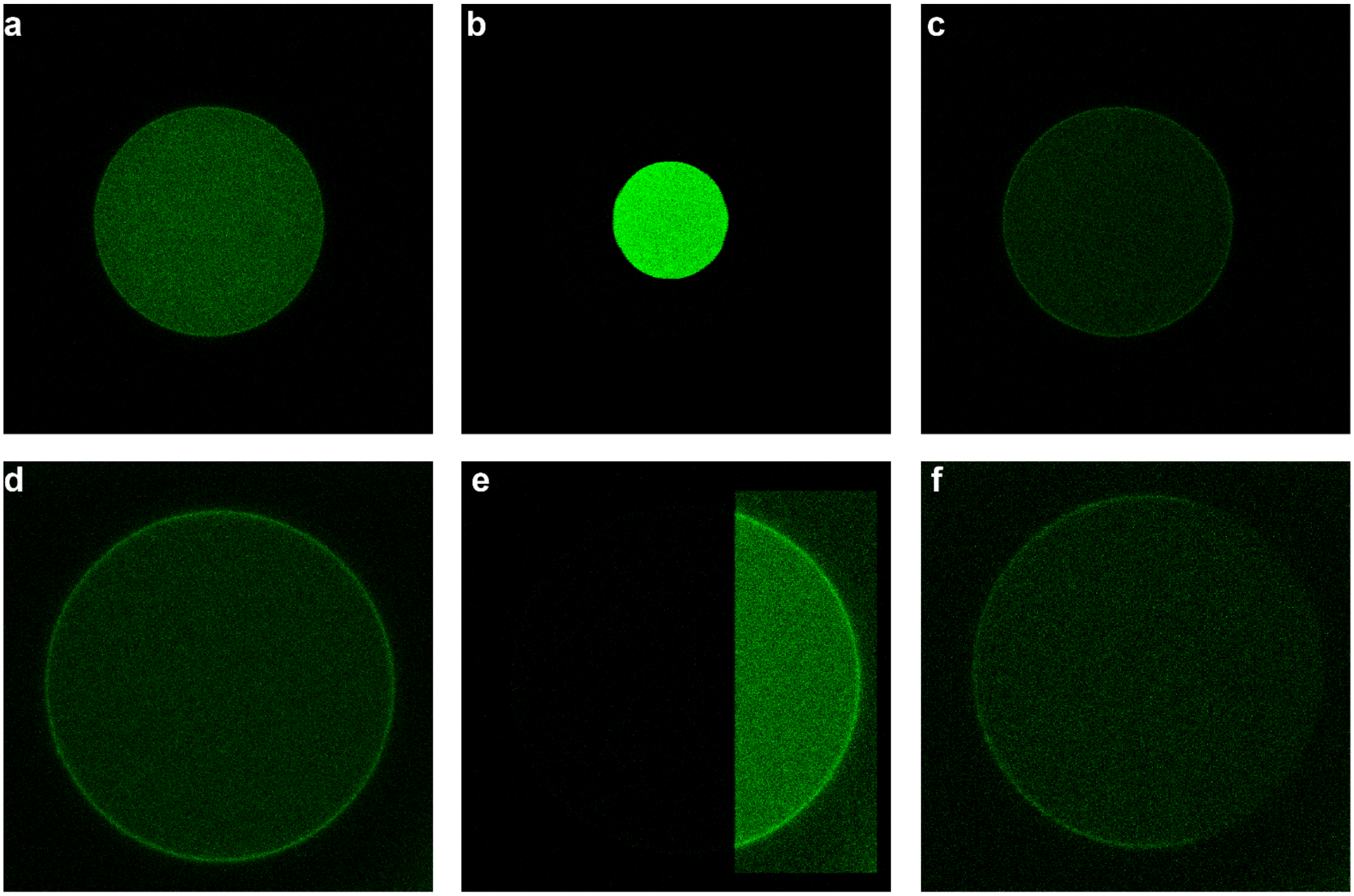


**Supplementary Fig. 4 | FRAP experiments on FITC-labeled proteinosomes containing streptavidin and free BSA-$NH_2$/PNIPAAm nanoconjugates. a,** Confocal micrograph of a high-**P** proteinosome before photo bleaching showing uniform green fluorescence **b**, Confocal micrograph of the circular bleach point in the middle of the proteinosome. **c**, Confocal micrograph of the proteinosome in **a** immediately after photobleaching for 1 minute using bleach point shown in **b**. The fluorescence in the interior of the proteinosome has been uniformly reduced indicating that the diffusion of the internalized FITC labeled conjugate is fast compared to the timescale of the bleaching process (1 minute). In contrast, fluorescence associated with the cross-linked BSA-$NH_2$/PNIPAAm membrane is more clearly visible. **d**, Confocal micrograph of a high-**P** proteinosome with a clearly visible membrane due to photo-bleaching of the interior. **e**, Confocal micrograph of a rectangular bleach point on the right side of the proteinosome shown in **d**. **f**, Confocal micrograph of the proteinosome in **d** after being photobleached for 1 minute using bleach point shown in **e** showing no recovery of the cross-linked FITC-labelled membrane.

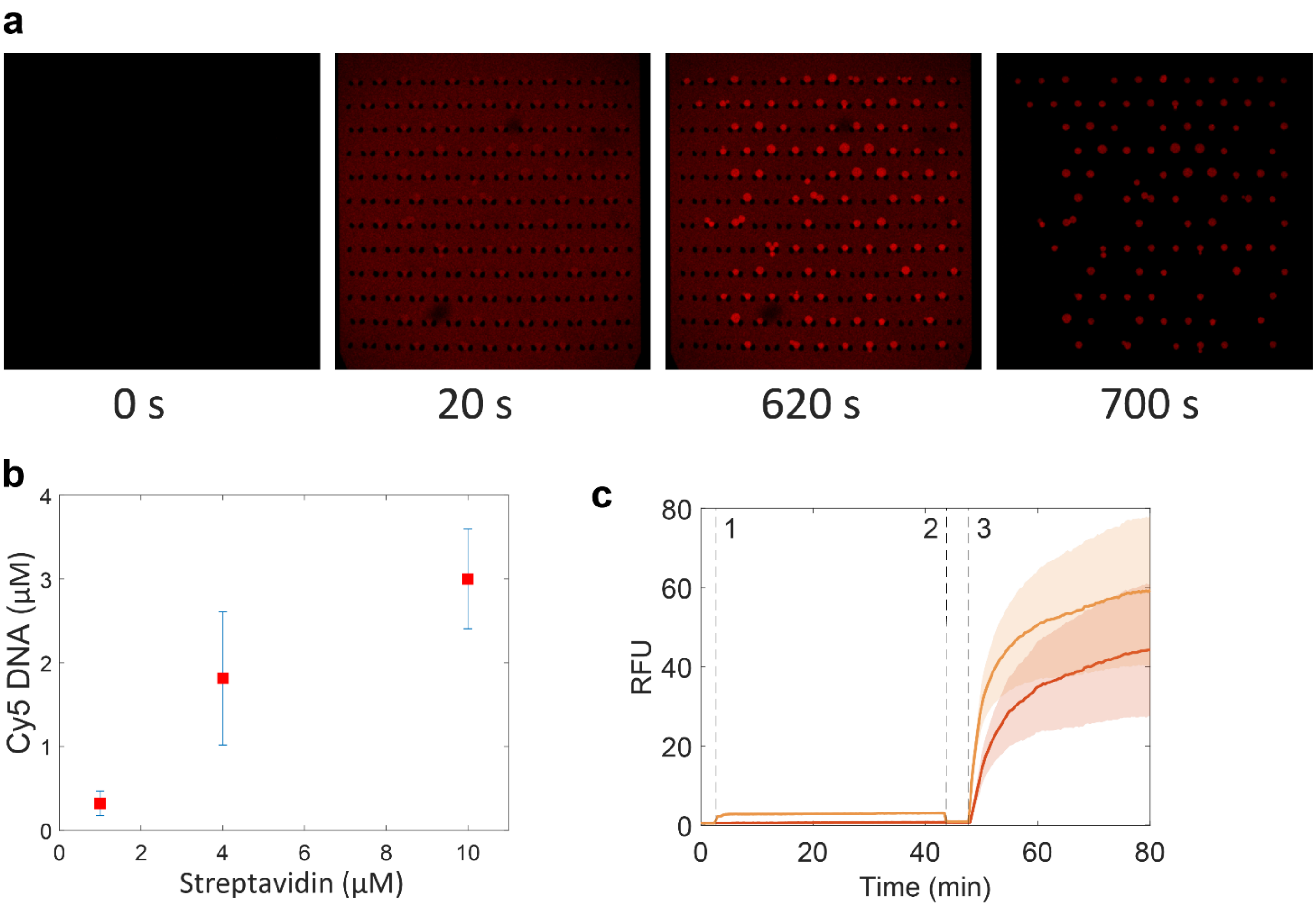


**Supplementary Fig. 5 | Anchoring of biotinylated ssDNA to proteinosome-encapsulated streptavidin and activating the anchored DNA gate complex. a,** Diffusion and localization of biotinylated DNA into streptavidin-containing high-**P** proteinosomes pre-organized within a microfluidic trapping device. At t=0 s, 1 μM of Cy5 labeled biotinylated ssDNA ($F_1$, Supplementary Table 1) is flowed into the microfluidic trapping chamber comprising an array of trapped streptavidin-containing proteinosomes. The labeled DNA strands immediately accumulate inside the proteinosomes by non-covalent binding to the encapsulated streptavidin. Saturated binding occurs within 620 s. DNA is then washed out of the chamber resulting in proteinosomes with entrapped *ss*DNA ($F_1$). In contrast, no localization is observed when non-biotinylated *ss*DNA is used (see Figure S1). In general, entrapment of *ss*DNA was undertaken in 1.5 mL Eppendorf tubes to produce larger quantities of the protocells and stored for later use at 4° C. **b**, A plot showing the DNA localization capacity of proteinosomes (Cy5-DNA concentration) prepared using a range of streptavidin concentrations. Concentrations are those used for the aqueous phase during the proteinosome assembly. **c**, Activating the entrapped DNA gate complex ($F_1$:$Q_1$ complex, Supplementary Table 1) **1,** addition of an input strand with a non-complementary toehold (A_NonComp_Cy3, 100 nM, Supplementary Table 1); **2,** washing to remove all the A_NonComp_Cy3, and **3,** addition of an input strand with a complementary toehold (A_Alexa546, 100 nM, Supplementary Table 1). Red trace is the Cy5 fluorescence of the activated DNA gate complex ($F_1$:A duplex). Orange trace is the Cy3/Alexa546 fluorescence of the input strand. The results show that the input with a non-complementary toehold can enter the proteinosomes (increase in orange trace at **1**), however it does not activate the DNA gate complex. In contrast, input strand with a complementary toehold can activate the DNA gate complex (increase in red trace at **3**). High-**P** proteinosomes prepared using 4 μM streptavidin were used in this experiment. This experiment was done a month after the DNA localization, demonstrating the long-term stability of the technique.

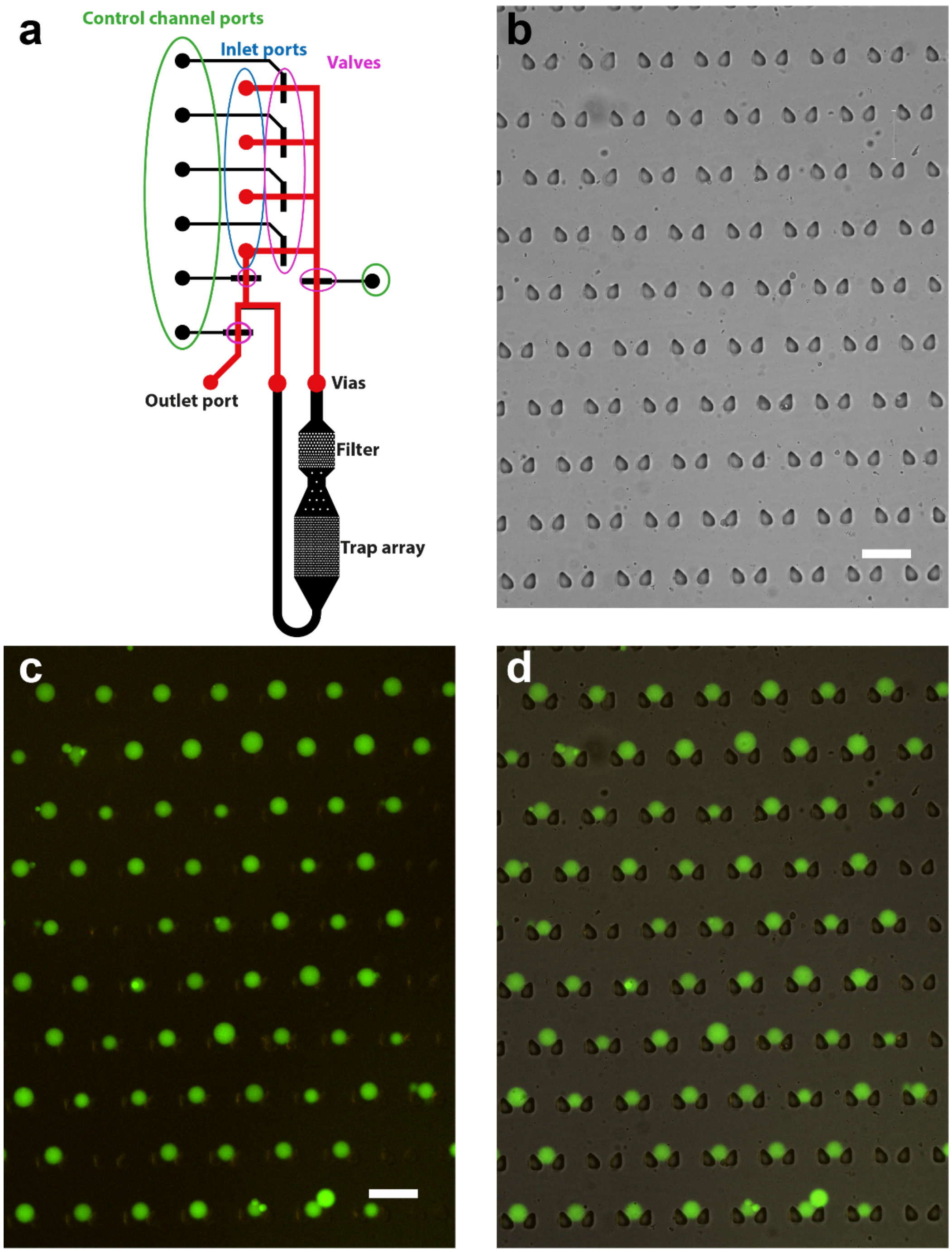


**Supplementary Fig. 6 | Design of the microfluidic device. a**, CAD drawing of the microfluidic device. Channels of the bottom layer (bonded to the glass slide) are in black. The rounded top layer channels are in red. The flow channels cross from the top layer to the bottom layer as only the rounded channels of the top layer can be closed using the pneumatic valves, while the trapping array requires the rectangular channels of the bottom layer. **b**, Bright-field micrograph of the trapping array. **c**, Epifluorescence micrograph of FITC labeled proteinosomes in the trapping array. **d**, Overlay of the epifluorescence image on the bright-field image. Scale bars are 100 µm.

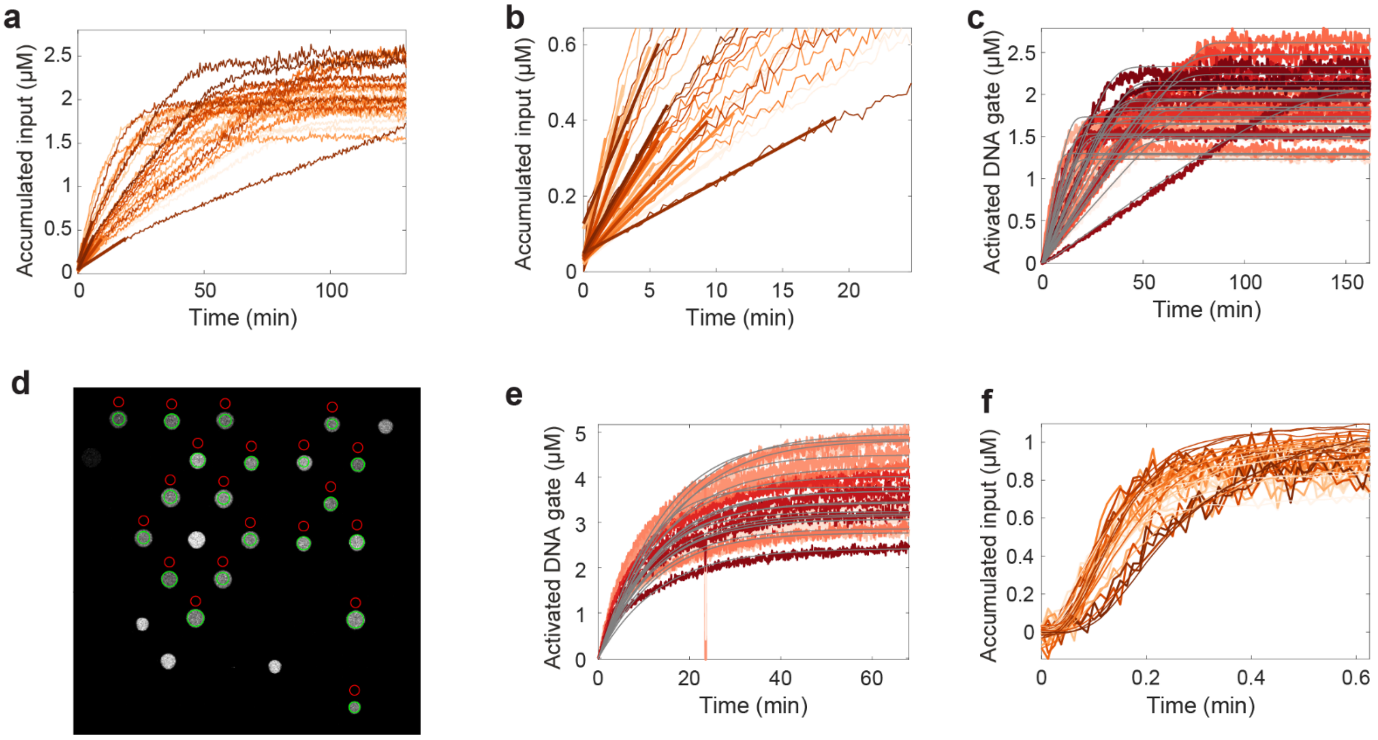


**Supplementary Fig. 7 | Parameter estimation of permeability coefficient (*P*) and DSD rate constant (*k*).** **a**, Experimental data showing the entry and accumulation of a fluorescently labeled input strand into a population of low-**P** proteinosomes. The experiment was initiated by flowing 1 µM of Alexa546 labeled input strand into the medium. The proteinosomes were prepared with 4 µM of streptavidin and contained the $\mathbf{F_1{:}Q_1}$ DNA complex. The DNA sequences are given in Supplementary Table 1. **b**, Close-up view of the initial 20 minutes of **a**, showing the fitted straight lines to estimate the initial slopes of the traces. As the diffusion is the rate limiting step for the low-***P*** proteinosomes, ***P*** could be calculated from the initial slopes using $P = \frac{\Delta[A]}{\Delta t}\frac{r}{3[A]_{\mathrm{out}}}$. **c**, Experimental data showing the activation of the fluorescent DNA gate complex and simulation results with the estimated DSD rate constant ***k*** and ***P***. A selection of this data is shown in Fig. 1g. **d**, A confocal micrograph of a population of high-***P*** proteinosomes used for estimating the parameters ***P*** and ***k***. Green circles indicate the proteinosomes that were used for analysis, the unmarked proteinosomes were excluded because they either moved during the experiment or exhibited highly abnormal reaction behavior. The red circles indicate the region from where the outside concentration of the labeled input strand was determined. **e**, Experimental data showing the activation of the fluorescent DNA gate in the population shown in **d** and simulation results with the estimated parameters ***k*** and ***P*.** A selection of this data is shown on Fig. 1g. The proteinosomes were prepared with 4 µM of streptavidin and contained the $\mathbf{F_1{:}Q_1}$ DNA complex. The reaction was triggered by flowing in 100 nM of the Alexa546 labeled input strand. **f**, Experimental data showing the entry of labeled input strand into the proteinosomes shown in **d**. This experiment was performed after the DSD reaction (**e**) had gone to completion and all free input had been washed away. 1 µM of the labeled input was used. This data was used to estimate parameter ***P***.

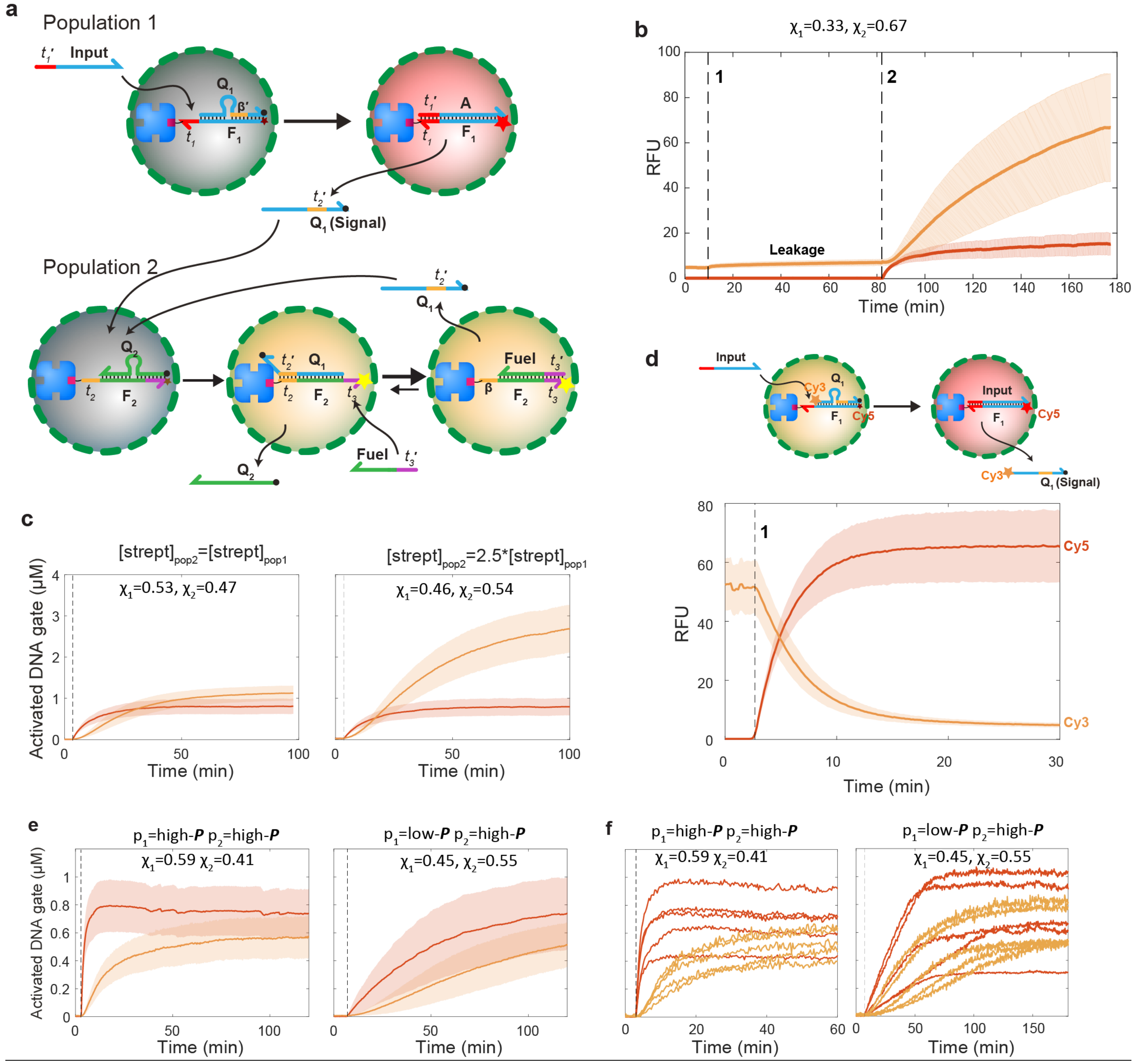


**Supplementary Fig. 8 | Design of the two-layer signaling cascade. a**, Molecular reaction diagram of the two-layer signaling cascade between two proteinosome populations. Coloration of graphics represents state of fluorescent probes: non-activated (grey), Cy5 activated (red), Alexa546 activated (orange). It is important to note that while for illustrative purposes the signal strand $\mathbf{Q}_1$ is shown to diffuse out of the population **2** proteinosome, it can also trigger the next reaction inside the same proteinosome by reacting with another inactive DNA gate complex. The sequences for this system are given in Supplementary Table 2. **b**, Leakage test of the two-layer cascade. When only fuel (1 µM) is injected into the external medium (time point 1), the protocells of the population **2** (orange profile) exhibit some activation due to leakage while population **1** (red profile) shows no response. The leakage reactions in DSD circuits are caused by impurities and misfolded DNA duplexes among other reasons and have been investigated in previous studies[2]; in this system, another possible cause for leakage are small amounts of free $Q_1$ strands that are not removed during the proteinosome washing step and also small amounts of proteinosome breakage during the loading procedure. At time point 2 (after 70 minutes), 100 nM of input strand and 1 µM of fuel strand is added to the system, resulting in the activation of population **1** (red curve) and, after a short delay population **2** responds (orange curve). The experiment was performed using high-***P*** proteinosomes prepared using 4 µM and 10 µM of streptavidin in populations **1** and **2,** respectively, with the fractions of the two populations, $\chi_1$=0.33, $\chi_1$=0.67. **c**, Comparison of the two-layer cascade with two different streptavidin concentrations in population **2** (left, 4 µM for both populations **1** and **2**; right, 4 µM for population **1,** 10 µM for population **2**). Experiments were initiated by injecting 100 nM of input and 1 µM of fuel strand into the external medium. **d**, Control experiment demonstrating the diffusion of the $\mathbf{Q}_1$ (signal) strand out of a single population of proteinosomes (population **1**; high-***P***, 4 µM of streptavidin). $\mathbf{Q_1}$ was labeled with a Cy3 fluorophore at the 5' end such that the proteinosomes exhibit high levels of Cy3 fluorescence before activation of the Cy5 fluorophore of the gate strand $\mathbf{F_1}$. The plots show time-dependent changes in the Cy5 and Cy3 fluorescence levels in the same population of proteinosomes. When input strand is added (time point 1), it displaces $\mathbf{Q_1}$ from $\mathbf{F_1}$**,** resulting in the increase of Cy5 fluorescence. At the same time, the Cy3 fluorescence decreases, confirming release of $\mathbf{Q_1}$ from $\mathbf{F_1}$ and its subsequent diffusion out of the proteinosomes. Experiments were initiated by injecting 100 nM of input strand into the external medium. Coloration of graphics; activated Cy3 (orange), activated Cy5 (red). **e**, Manipulating the community-level behavior of the binary population by tuning the permeability of the protocell membrane. With exclusively high-P protocells (left), both populations provide a fast response to the added input strand, swapping the first population to low-P significantly slows the response rates of both populations (right). Experiments were initiated by injecting 500 nM of input strand to the external medium, all proteinosome populations were assembled using 4 µM of streptavidin. **f**, Time dependent activation levels of five individual protocells from the populations shown in **e**, revealing a very clear difference in the response profiles of population **1** protocells between the high-***P*** and low-***P*** versions. With low-***P*** population **1**, the response of population **2**, protocells also becomes more linear, however this effect is limited as the concentration of $\mathbf{Q}_1$ that the population **2** protocells are subjected to is determined by the collective response of population **1**, where the linearity is lost due to polydispersity in response rates and final values.

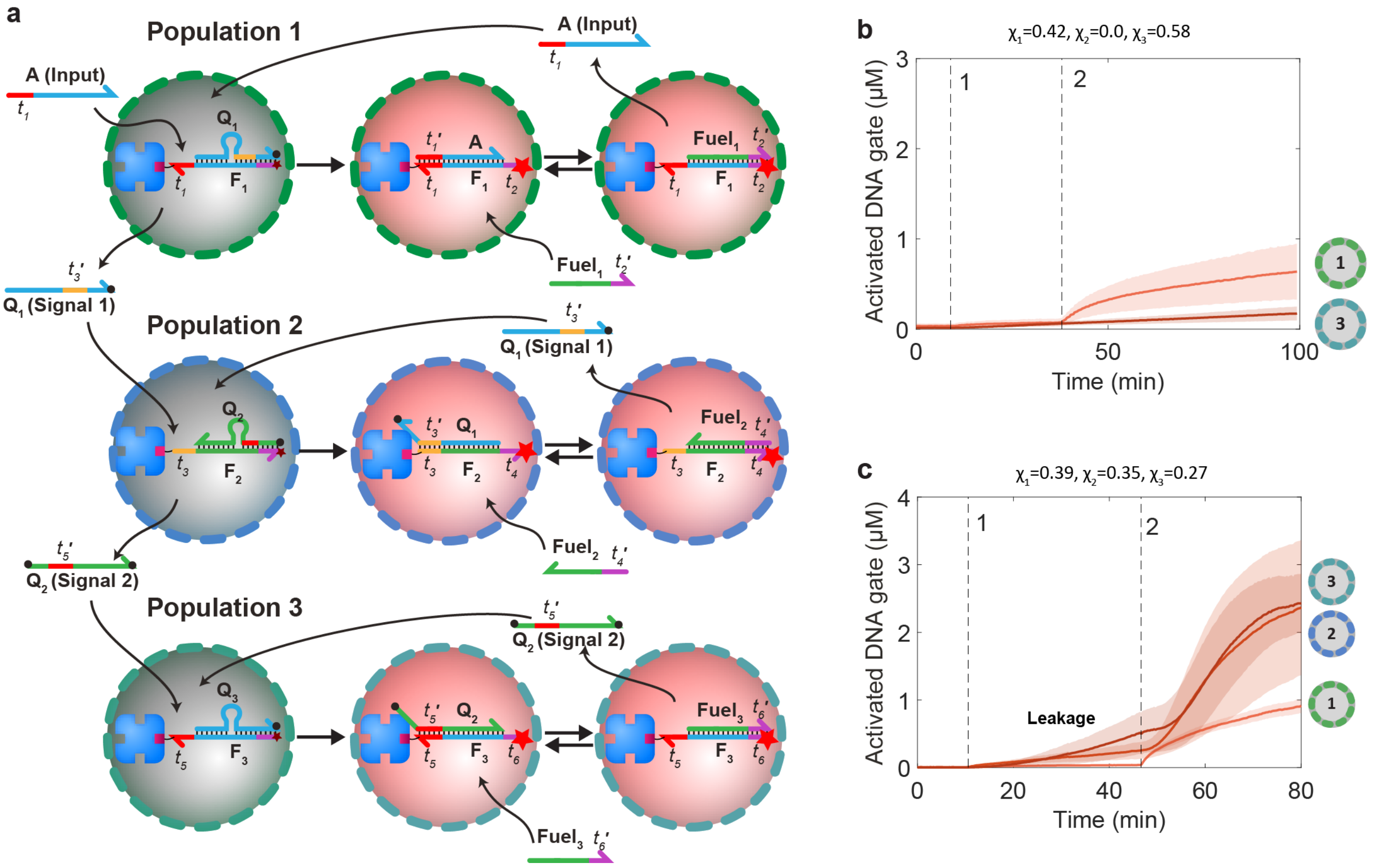


**Supplementary Fig. 9 | Design of the three-layer signaling cascade. a,** Molecular reaction diagram of the three-layer signaling cascade using three proteinosome populations in sequence order **1,2,3**. Coloration of graphics represents state of fluorescent probes: non-activated (grey), Cy5 activated (red). The DNA sequences for this system are given in Supplementary Table 3. **b**, Control experiment showing the response of the system consisting only of populations **1** and **3.** At time point 1, the three fuel strands were flowed into the protocell community followed by both fuel and input at time point 2. The reaction was performed using 10 nM of input and 500 nM of each of the three fuel strands. Both proteinosome populations were high-***P*** and were assembled with 4 µM of streptavidin. **c**. Leakage test of the three-layer cascade. At time point 1, the three fuel strands (500 nM each) were flowed into the protocell community. The leakage signal is amplified in every successive step with population **3** exhibiting the strongest signal. At time point 2, (approximately 40 minutes later), the fuels (500 nM each) and input strand (10 nM) are added to the community; protocells of all three populations produce their expected response with increasing delays. The successive amplification of the leakage signal means that in order to construct catalytic cascades with more stages, additional design considerations would need to be considered to keep the leakage signal under control. All proteinosome populations were high-***P*** and were assembled with 4 µM of streptavidin.

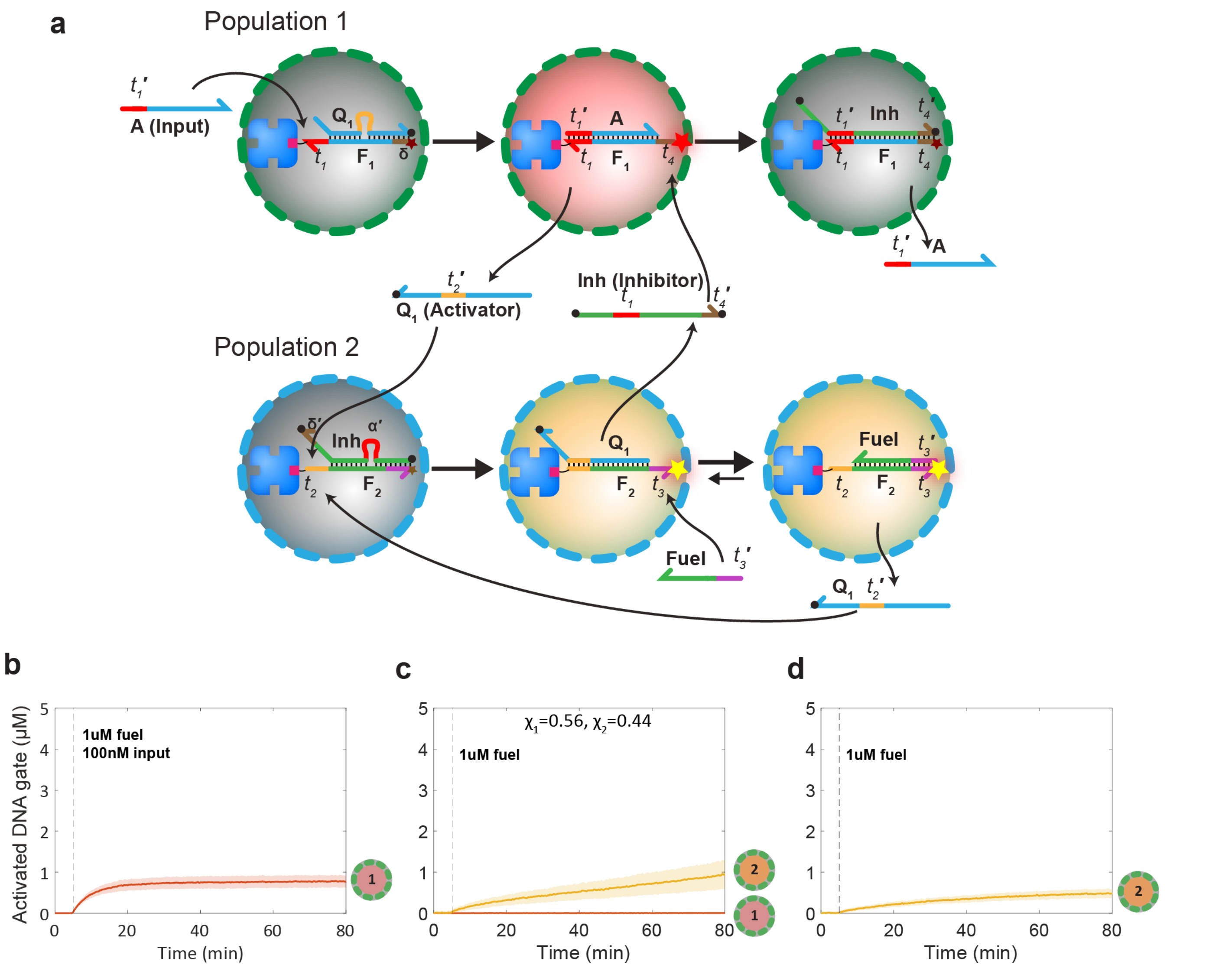


**Supplementary Fig. 10 | Design of the negative feedback network. a,** Molecular reaction diagram of the negative feedback network between two proteinosome populations. Coloration of graphics represents state of fluorescent probes: non-activated (grey), Cy5 activated (red), Alexa546 activated (orange). The sequences for this system are given in Supplementary Table 4. **b**, A control experiment with only population **1** (high-***P*** prepared using 4 μM of streptavidin) showing time-dependent changes in concentration (fluorescence) due to activation of the streptavidin-anchored DNA gate complex (**$F_1$:A**) in the absence of a released inhibitor strand. Reaction was performed with 200 nM of the input and 1 μM of the fuel strand; both were added at the time point indicated by the dashed line. **c**, Control experiment to evaluate the leakage reaction of the catalytic module in the absence of the input strand for a mixed community of protocell populations **1** and **2** ($\chi_1$=0.56, $\chi_2$=0.44; both high-***P*** prepared using 4 μM and 10 μM of streptavidin in population **1** and **2** respectively). Reaction was performed with 1 μM of fuel. **d**, Control experiment to evaluate the leakage reaction of the catalytic module for population **2** alone in the absence of the input strand (high-***P***, 10 μM of streptavidin). Reaction was performed with 1 μM of the fuel strand. The results from **c** and **d** reveal that the total leakage is a combination of the inherent leakage of the catalytic module in population **2** (**d**) and the presence of free **$Q_1$** in population **1**.

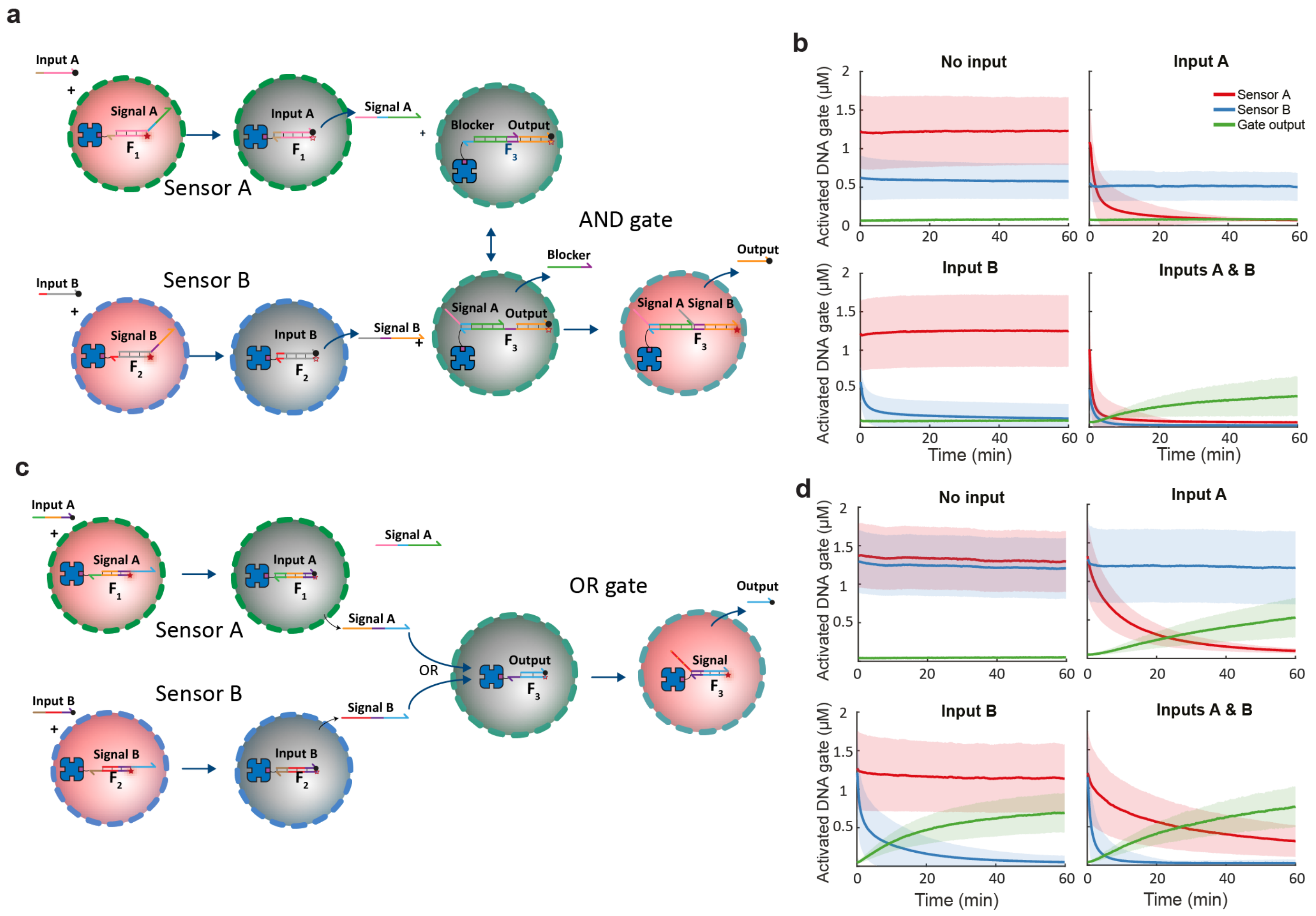


**Supplementary Fig. 11 | Design of the combinatorial sensing-and-processing devices for distributed computing in protocell communities. a**, Molecular reaction diagram of the protocell-based three-population AND network based on the synergistic activity of two transducers and a single receiver. The sensor proteinosomes (populations **A** and **B**) contain DNA gate complexes that initially consist of a streptavidin-anchored fluorescently active strand (**$F_1$** or **$F_2$**) and signalling strands (**A** or **B**), respectively. **$F_1$** and **$F_2$** become quenched upon binding of the cognate input strands, which in turn releases signaling strands **A** and **B** from the two different sensor populations. The population of AND gate proteinosomes contain a DNA gate complex consisting of a deactivated strand (**$F_3$**) bound to two complementary oligonucleotides that function either as a blocker or quencher/output strand during processing. Activation of the complex requires the binding of both the signal strands such that **$F_3$** acts as a fluorescent probe of the AND operation. Signal **A** binds to the initially free toehold of **$F_3$**, displaces the blocker strand and thereby reveals a second toehold where signal **B** can bind and displace the output (quencher) strand. Non-activated protocells (grey); Cy5-activated protocells (red). The DNA sequences for this system are given in Supplementary Table 5. **b**, Experimental data for the three-population AND network with all four possible input configurations. Profiles are shown for DNA gate complex activation levels of **$F_1$** (red), **$F_2$** (blue) and **$F_3$** (green). The results demonstrate that the sensor proteinosomes produce a response when the appropriate input strand is present. The AND gate proteinosomes only produce a response when both sensor populations have been activated. All three populations were high-***P*** proteinosomes prepared using 4 µM of streptavidin; fractions of the population were within the range of $\chi_A$=0.40-0.49, $\chi_B$=0.41-0.48, $\chi_{AND}$=0.06-0.16. The concentrations of the input strands (**A** and **B**) when present were 100 nM. **c**, Molecular reaction diagram of the protocell-based three-population OR network based on the equivalence in activity of two transducers and a single receiver. The design concept is similar to that shown in **a** but with different input, gate and signal strands in the two fluorescent sensor protocell populations, and with a population of OR gate proteinosomes containing a deactivated gate complex that can be fluorescently activated by signal strands originating from either upstream sensor population. Non-activated protocells (grey); Cy5-activated protocells (red). **d**, Experimental data for the three-population OR network with all four possible input configurations. Profiles are shown for DNA gate complex activation levels of $\mathbf{F_1}$ (red), $\mathbf{F_2}$ (blue) and $\mathbf{F_3}$ (green).The results demonstrate that the OR gate proteinosomes produce a response when at least one of the sensor populations have been activated. All three populations were high-***P*** proteinosomes prepared with 4 μM of streptavidin; fractions of the ternary population were $\chi_A$=0.37-0.47, $\chi_B$=0.45-0.54, $\chi_{AND}$=0.08-0.15. The concentrations of the input strands (**A** and **B**), when present, were 100 nM.

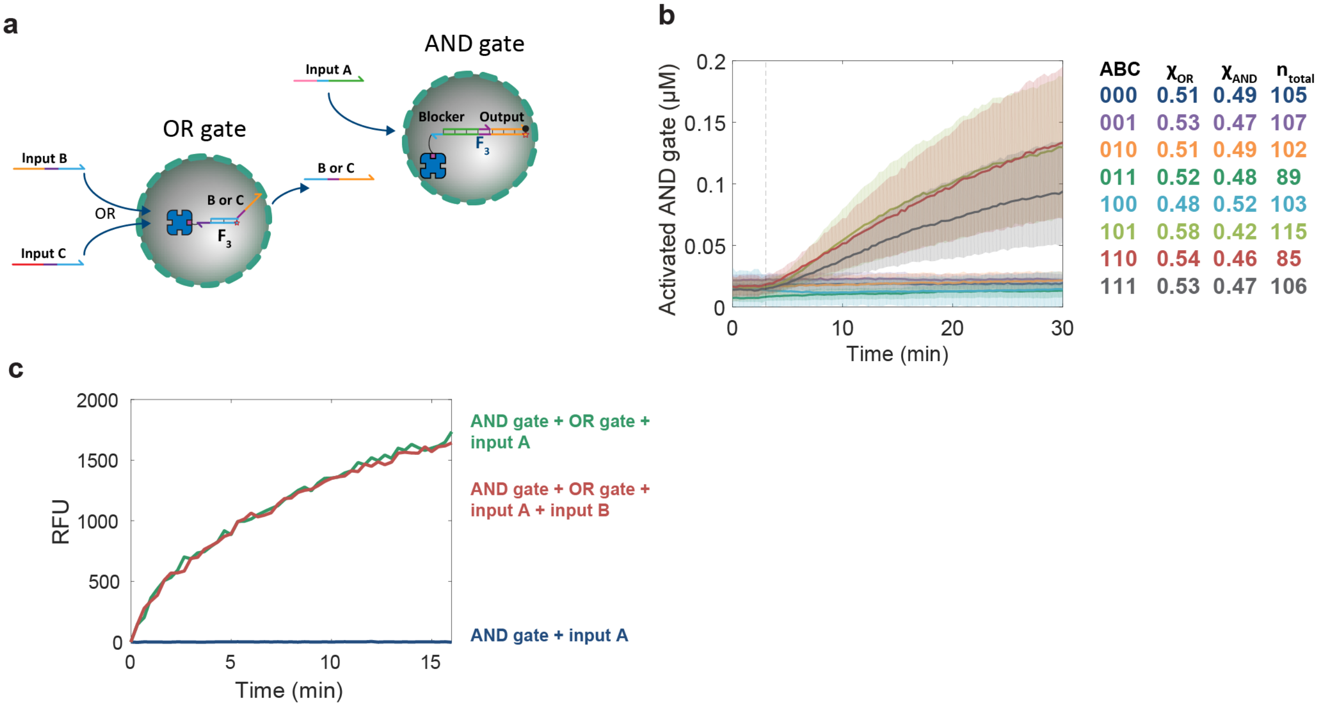


**Supplementary Fig. 12 | Design of the serial logic circuit. a**, Reaction diagram of the serial logic circuit. The system is constructed from the individual gate circuits shown in Supplementary Figure 11a and c. Specifically, a deactivated AND module (**$F_3$**:blocker:quencher/output complex) capable of sensing DNA signalling strands **A** and **B** is combined with a deactivated OR module (**$F_3$**:output complex), which can sense input strands **B** and **C.** Critically, the output strand of the OR gate is extended with the sequence corresponding to signal **B** of the AND gate. As a consequence, inputs of **B** or **C** into the OR module produce an output strand that serves as an input for the AND gate. In the presence of input strand **A**, this leads to activation of the AND gate protocells and a fluorescence read-out due to displacement of the quencher strand as shown in Supplementary Figure 11a. **b**, Experimental data of the two-gate, three input logic circuit with means and standard deviations of all eight input configurations. The color-coded traces correspond to the rows of the table on the right which indicates the specific input configuration (presence (1) or absence (0) of the three input strands **A**, **B** and **C**), fractions of the two populations and the total number of proteinosomes for the corresponding experiment. The experiments were performed with high-***P*** proteinosomes prepared with 4 µM of streptavidin. The concentration of each input strand, when present, was 500 nM. **c**, Baseline corrected fluorescence traces of batch experiments with the serial logic circuit. The final concentrations of all strands (AND gate complex, OR gate complex, input A and input B) were 100 nM. The components, except input **A**, were mixed in a reaction buffer and the Cy5-fluorescence (RFU = relative fluorescence units) was measured on a plate reader and the baseline was recorded. At t=0 the measurement was paused, Input **A** was added into all wells and the measurement was resumed. The results show that the presence of inactive OR gate yields a similar level of AND gate activation as the activated OR gate indicating that under batch conditions the two modules of the serial logic circuit cross-hybridize and thereby render it inoperative. The sequences for this system are given in Supplementary Table 5.

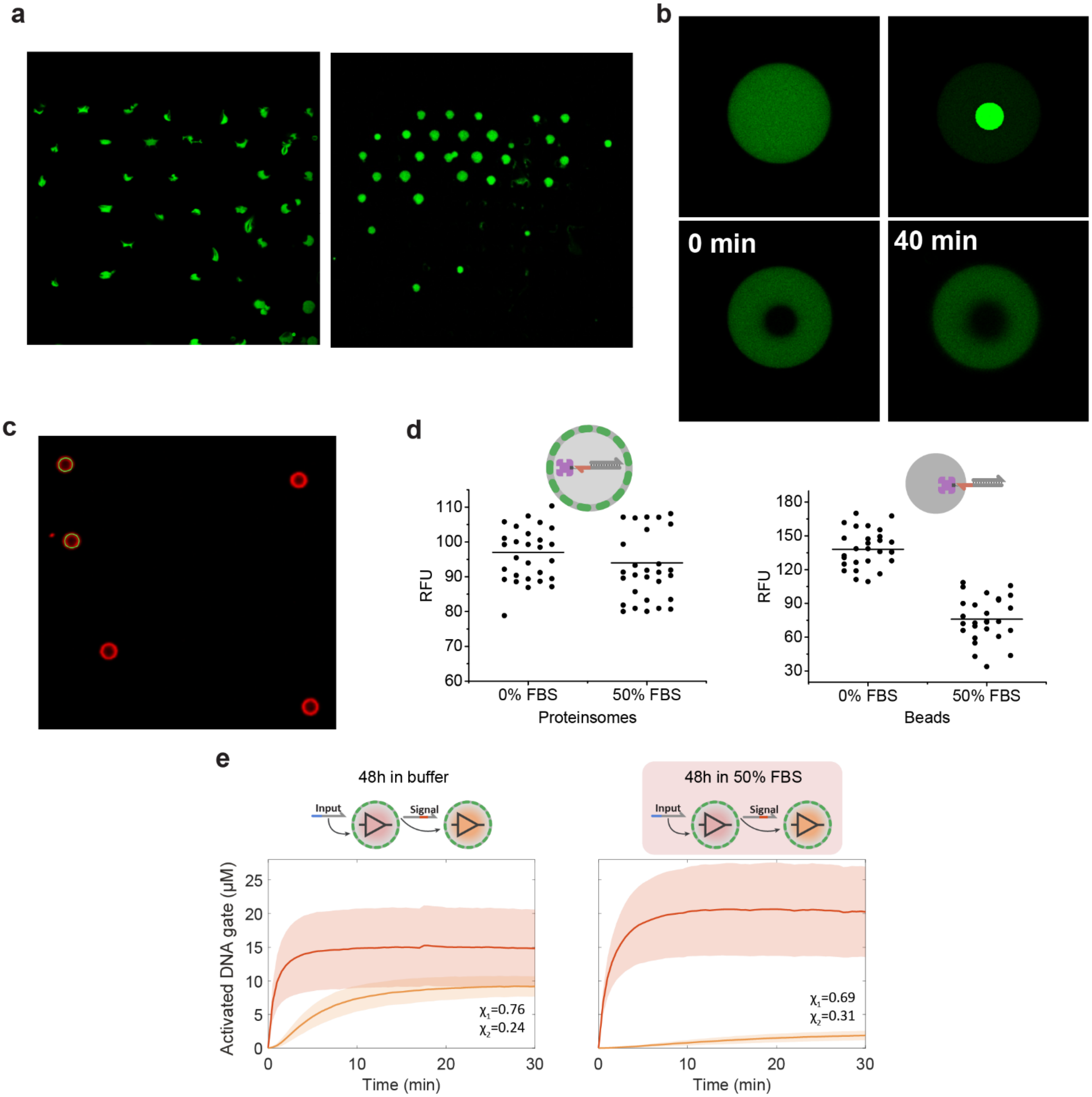


**Supplementary Fig. 13 | Proteinosomes and DSD circuitry in 50% FBS. a**, Confocal micrographs of microfluidic trapped arrays of high-***P*** proteinosomes prepared without (left side) or with (right side) encapsulated non-functionalized BSA (60 mg/mL) and recorded immediately after filling the trapping chamber with 50% FBS. When exposed to FBS the proteinosomes prepared without additional BSA collapsed immediately, while the proteinosomes with additional BSA retained their morphology. The proteinosomes were prepared using 20 µM of streptavidin. **b**, FRAP experiments elucidating the structure of the proteinosomes that were assembled with 60 mg/mL non-functionalized BSA. The top two micrographs show a proteinosome before photo-bleaching and the bleach-point, the bottom micrographs show the proteinosome immediately after and 40 minutes after photo-bleaching. The interior is much more gel-like with only minimal amount of signal recovery in the center of the proteinosome 40 minutes after photo-bleaching. **c**, A confocal micrograph of streptavidin-coated polystyrene beads, functionalized with Cy5-labeled DNA. The mean RFU value of the beads was calculated by averaging the RFU values over a circular path, as illustrated by the two green circles. **d**, Final values of DSD reactions performed in 50% FBS in proteinosomes and on streptavidin coated beads. The experiments with beads were performed by localizing the **F:Q** complex (Supplementary Methods and Supplementary Table 6) on the surface of the beads (approximately $8.4 \times 10^{-11}$ umol, per bead), adding the input DNA (2 µM) and measuring the final fluorescence values using confocal microscope. The proteinosomes were prepared with 20 µM streptavidin and 60 mg/mL BSA (Methods). 2 µM of input DNA was used to trigger the DSD reaction in proteinosomes. The data acquisition methodology for beads is described in Supplementary Methods. The results show that after 48h incubation in 50% FBS, on an average the **F:Q** gate complex retains 55% of its activity on beads and 97% in proteinosomes. **e**, Comparison of the two-layer transducer-receiver signaling cascade (Fig. 4e) in buffer and in 50% FBS. In the first case, the proteinosomes of the two populations were mixed together and incubated for 48h (Methods) at room temperature in a buffer solution, localized in the trapping chamber and then the cascade was activated by adding 2 µM of input DNA. In the second case all the steps were performed in 50% FBS. The results show that the response of the transducer population does not change significantly, while the response of the receiver population is much weaker due to signal strand degradation in FBS.

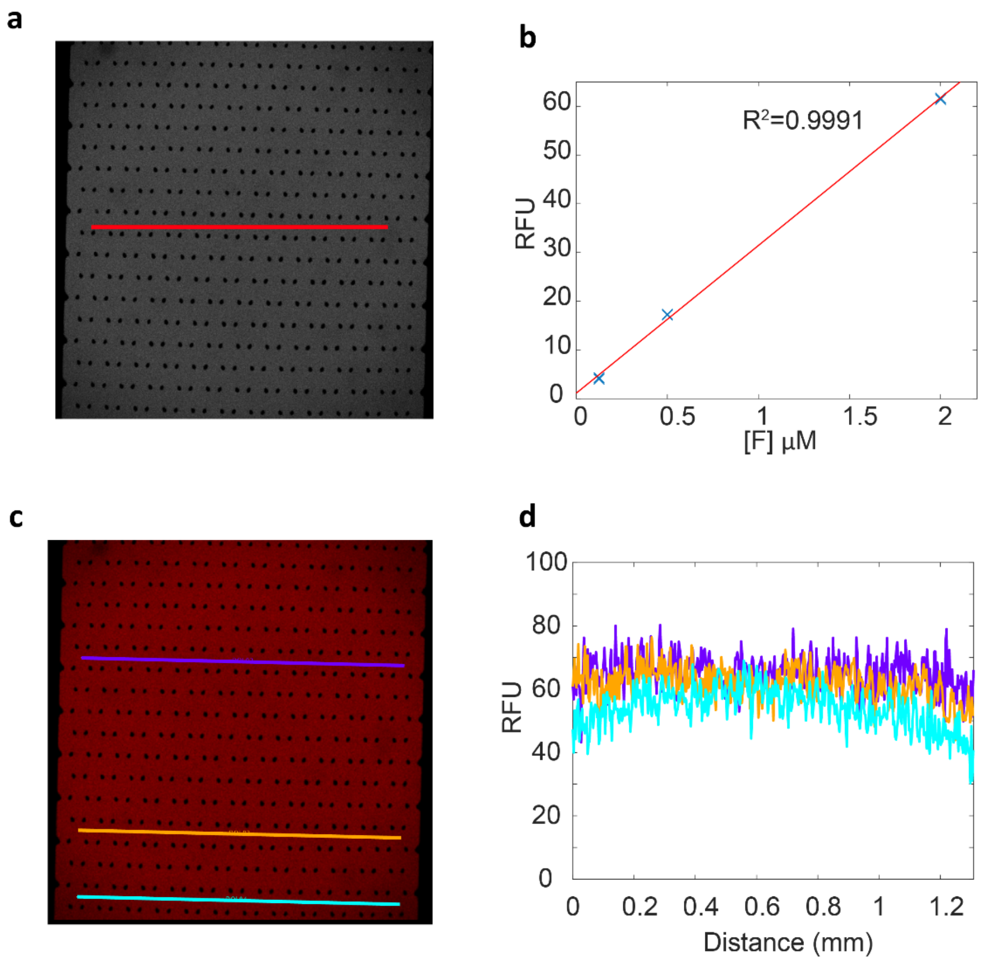


**Supplementary Fig. 14 | Verification of the linearity of the concentration estimation method and evaluation of the spatial distortion in confocal measurements. a**, A confocal micrograph of the microfluidic trapping device filled fluorescently labeled DNA. The RFU-to-concentration conversion factor for a specific DNA gate complex was obtained by filling the device with the active form of the gate complex at 1 μM and measuring the average RFU value across a straight horizontal line through the device. **b**, The linearity of the conversion method was verified by performing the measurement shown in **a** over a range of activated DNA gate complex concentrations (100 nM, 500 nM and 2 μM), fitting a straight line through the data and calculating the coefficient of determination ($R^2$). **c**, To verify that there is no distortion of the fluorescence measurements depending on the position in the device, the fluorescence readings were compared for different positions in the device. **d**, Position vs fluorescence intensity profiles across the straight lines shown in **c**. Noticeable distortion can only be seen in the corners of the image (ends of the cyan line).

### Supplementary Notes

**Supplementary note 1 | Derivation of the 1D model.**

Assuming a perfectly spherical proteinosome, we can rely on its symmetry and describe the diffusion through its membrane using Fick's first law in one dimension

$$\text{Eq. 1} \quad J_i = -D\frac{\partial c_i}{\partial x}$$

where $J_i$ is the flux of species *i* per unit area per unit time (mol $m^{-2}$ $s^{-1}$), $D$ is the diffusion coefficient ($m^2$ $s^{-1}$), $c_i$ is the concentration (mol $m^{-3}$) and $x$ is the position (m). Looking at diffusion just through the membrane and assuming that both inside and outside the system is well mixed we can write

$$\text{Eq. 2} \quad J_i = -D\frac{C_{\mathrm{in}} - C_{\mathrm{out}}}{\Delta x} = P(C_{\mathrm{out}} - C_{\mathrm{in}})$$

where $\Delta x$ is the thickness of the membrane, $P = \frac{D}{\Delta x}$ is the permeability coefficient of the membrane (m $s^{-1}$). Now $J_i$ is the flux per unit area into the protocell, the rate of change of the number of molecules inside the protocell is then simply

$$\text{Eq. 3} \quad \frac{dn_i}{dt} = AJ_i$$

where $A$ is the area of the membrane. Dividing by the volume gives the change in concentration

$$\text{Eq. 4} \quad \frac{dc_{in}}{dt} = \frac{A}{V}J_i = \frac{3}{r}P(c_{\mathrm{out}} - c_{in})$$

where $r$ is the radius and $V$ is the volume of the protocell. This is equivalent to Eq. 1 of main text and was used to estimate the permeability coefficient of the proteinosomes.

**Supplementary note 2 | Visual DSD code of the 2D simulations of the 2-layer cascade.**

```
directive simulator pde
directive simulation {
  final= 100.000000; //min
  points= 100;
  plots=[ F1Act ; F2Q1 ; F2Fuel; ActOut]
}

directive spatial {
  dimensions = 2;
  dt = 0.020000; //min
  xmax = 1500; //um
  nx = 512;
  boundary = ZeroFlux;
  diffusibles = [Q1sol = 2227.458030 ; ActOut = 2227.458030 ; FuelOut = 2669.276407 ; Q2sol =
2227.458030];  //um^2 min^-1
}

directive parameters [
  rprot = 20; //um
  kdis = 0.000930;  //nm^-1 min^-1
  kFuel = 0.000930; //nm^-1 min^-1
  P = 85.000000; //um min^-1
]

| init psA {spatial = { points = [
  { width = 0.026667; value = 1; x = 0.020000; y = 0.020000 };
  { width = 0.026667; value = 1; x = 0.260000; y = 0.020000 };
  { width = 0.026667; value = 1; x = 0.580000; y = 0.020000 };
  { width = 0.026667; value = 1; x = 0.660000; y = 0.020000 };
  { width = 0.026667; value = 1; x = 0.820000; y = 0.020000 };
  { width = 0.026667; value = 1; x = 0.980000; y = 0.020000 };
  { width = 0.026667; value = 1; x = 0.300000; y = 0.100000 };
  { width = 0.026667; value = 1; x = 0.540000; y = 0.100000 };
  { width = 0.026667; value = 1; x = 0.620000; y = 0.100000 };
  { width = 0.026667; value = 1; x = 0.700000; y = 0.100000 };
  { width = 0.026667; value = 1; x = 0.940000; y = 0.100000 };
  { width = 0.026667; value = 1; x = 0.020000; y = 0.180000 };
  { width = 0.026667; value = 1; x = 0.180000; y = 0.180000 };
  { width = 0.026667; value = 1; x = 0.260000; y = 0.180000 };
  { width = 0.026667; value = 1; x = 0.340000; y = 0.180000 };
  { width = 0.026667; value = 1; x = 0.420000; y = 0.180000 };
  { width = 0.026667; value = 1; x = 0.820000; y = 0.180000 };
  { width = 0.026667; value = 1; x = 0.980000; y = 0.180000 };
  { width = 0.026667; value = 1; x = 0.060000; y = 0.260000 };
  { width = 0.026667; value = 1; x = 0.140000; y = 0.260000 };
  { width = 0.026667; value = 1; x = 0.220000; y = 0.260000 };
  { width = 0.026667; value = 1; x = 0.460000; y = 0.260000 };
  { width = 0.026667; value = 1; x = 0.700000; y = 0.260000 };
  { width = 0.026667; value = 1; x = 0.020000; y = 0.340000 };
  { width = 0.026667; value = 1; x = 0.180000; y = 0.340000 };
  { width = 0.026667; value = 1; x = 0.260000; y = 0.340000 };
  { width = 0.026667; value = 1; x = 0.340000; y = 0.340000 };
  { width = 0.026667; value = 1; x = 0.500000; y = 0.340000 };
  { width = 0.026667; value = 1; x = 0.580000; y = 0.340000 };
  { width = 0.026667; value = 1; x = 0.740000; y = 0.340000 };
  { width = 0.026667; value = 1; x = 0.820000; y = 0.340000 };
  { width = 0.026667; value = 1; x = 0.980000; y = 0.340000 };
  { width = 0.026667; value = 1; x = 0.060000; y = 0.420000 };
  { width = 0.026667; value = 1; x = 0.140000; y = 0.420000 };
  { width = 0.026667; value = 1; x = 0.220000; y = 0.420000 };
  { width = 0.026667; value = 1; x = 0.300000; y = 0.420000 };
  { width = 0.026667; value = 1; x = 0.380000; y = 0.420000 };
  { width = 0.026667; value = 1; x = 0.460000; y = 0.420000 };
  { width = 0.026667; value = 1; x = 0.540000; y = 0.420000 };
  { width = 0.026667; value = 1; x = 0.700000; y = 0.420000 };
  { width = 0.026667; value = 1; x = 0.020000; y = 0.500000 };
  { width = 0.026667; value = 1; x = 0.100000; y = 0.500000 };
  { width = 0.026667; value = 1; x = 0.180000; y = 0.500000 };
  { width = 0.026667; value = 1; x = 0.260000; y = 0.500000 };
  { width = 0.026667; value = 1; x = 0.340000; y = 0.500000 };
  { width = 0.026667; value = 1; x = 0.500000; y = 0.500000 };
  { width = 0.026667; value = 1; x = 0.660000; y = 0.500000 };
```

```
  { width = 0.026667; value = 1; x = 0.740000; y = 0.500000 };
  { width = 0.026667; value = 1; x = 0.900000; y = 0.500000 };
  { width = 0.026667; value = 1; x = 0.980000; y = 0.500000 };
  { width = 0.026667; value = 1; x = 0.060000; y = 0.580000 };
  { width = 0.026667; value = 1; x = 0.220000; y = 0.580000 };
  { width = 0.026667; value = 1; x = 0.300000; y = 0.580000 };
  { width = 0.026667; value = 1; x = 0.460000; y = 0.580000 };
  { width = 0.026667; value = 1; x = 0.540000; y = 0.580000 };
  { width = 0.026667; value = 1; x = 0.700000; y = 0.580000 };
  { width = 0.026667; value = 1; x = 0.860000; y = 0.580000 };
  { width = 0.026667; value = 1; x = 0.020000; y = 0.660000 };
  { width = 0.026667; value = 1; x = 0.260000; y = 0.660000 };
  { width = 0.026667; value = 1; x = 0.340000; y = 0.660000 };
  { width = 0.026667; value = 1; x = 0.420000; y = 0.660000 };
  { width = 0.026667; value = 1; x = 0.660000; y = 0.660000 };
  { width = 0.026667; value = 1; x = 0.740000; y = 0.660000 };
  { width = 0.026667; value = 1; x = 0.060000; y = 0.740000 };
  { width = 0.026667; value = 1; x = 0.220000; y = 0.740000 };
  { width = 0.026667; value = 1; x = 0.380000; y = 0.740000 };
  { width = 0.026667; value = 1; x = 0.460000; y = 0.740000 };
  { width = 0.026667; value = 1; x = 0.540000; y = 0.740000 };
  { width = 0.026667; value = 1; x = 0.620000; y = 0.740000 };
  { width = 0.026667; value = 1; x = 0.700000; y = 0.740000 };
  { width = 0.026667; value = 1; x = 0.780000; y = 0.740000 };
  { width = 0.026667; value = 1; x = 0.860000; y = 0.740000 };
  { width = 0.026667; value = 1; x = 0.020000; y = 0.820000 };
  { width = 0.026667; value = 1; x = 0.180000; y = 0.820000 };
  { width = 0.026667; value = 1; x = 0.340000; y = 0.820000 };
  { width = 0.026667; value = 1; x = 0.500000; y = 0.820000 };
  { width = 0.026667; value = 1; x = 0.580000; y = 0.820000 };
  { width = 0.026667; value = 1; x = 0.060000; y = 0.900000 };
  { width = 0.026667; value = 1; x = 0.300000; y = 0.900000 };
  { width = 0.026667; value = 1; x = 0.380000; y = 0.900000 };
  { width = 0.026667; value = 1; x = 0.460000; y = 0.900000 };
  { width = 0.026667; value = 1; x = 0.620000; y = 0.900000 };
  { width = 0.026667; value = 1; x = 0.700000; y = 0.900000 };
  { width = 0.026667; value = 1; x = 0.780000; y = 0.900000 };
  { width = 0.026667; value = 1; x = 0.860000; y = 0.900000 };
  { width = 0.026667; value = 1; x = 0.100000; y = 0.980000 };
  { width = 0.026667; value = 1; x = 0.260000; y = 0.980000 };
  { width = 0.026667; value = 1; x = 0.420000; y = 0.980000 };
  { width = 0.026667; value = 1; x = 0.900000; y = 0.980000 };
  { width = 0.026667; value = 1; x = 0.980000; y = 0.980000 };
  ] } }

| init F1Q1 {spatial = { points = [
  { width = 0.026667; value = 3845.990707; x = 0.020000; y = 0.020000 };
  { width = 0.026667; value = 2654.710600; x = 0.260000; y = 0.020000 };
  { width = 0.026667; value = 4579.186852; x = 0.580000; y = 0.020000 };
  { width = 0.026667; value = 3497.047867; x = 0.660000; y = 0.020000 };
  { width = 0.026667; value = 4007.967186; x = 0.820000; y = 0.020000 };
  { width = 0.026667; value = 3417.814158; x = 0.980000; y = 0.020000 };
  { width = 0.026667; value = 2935.440745; x = 0.300000; y = 0.100000 };
  { width = 0.026667; value = 2846.192882; x = 0.540000; y = 0.100000 };
  { width = 0.026667; value = 3877.204587; x = 0.620000; y = 0.100000 };
  { width = 0.026667; value = 3774.479469; x = 0.700000; y = 0.100000 };
  { width = 0.026667; value = 3918.013558; x = 0.940000; y = 0.100000 };
  { width = 0.026667; value = 3265.107910; x = 0.020000; y = 0.180000 };
  { width = 0.026667; value = 3509.211403; x = 0.180000; y = 0.180000 };
  { width = 0.026667; value = 3682.571024; x = 0.260000; y = 0.180000 };
  { width = 0.026667; value = 2414.018663; x = 0.340000; y = 0.180000 };
  { width = 0.026667; value = 4540.633176; x = 0.420000; y = 0.180000 };
  { width = 0.026667; value = 3073.924565; x = 0.820000; y = 0.180000 };
  { width = 0.026667; value = 3815.312505; x = 0.980000; y = 0.180000 };
  { width = 0.026667; value = 2793.333455; x = 0.060000; y = 0.260000 };
  { width = 0.026667; value = 3941.093972; x = 0.140000; y = 0.260000 };
  { width = 0.026667; value = 3793.585412; x = 0.220000; y = 0.260000 };
  { width = 0.026667; value = 2946.220470; x = 0.460000; y = 0.260000 };
  { width = 0.026667; value = 4005.216900; x = 0.700000; y = 0.260000 };
  { width = 0.026667; value = 2623.727184; x = 0.020000; y = 0.340000 };
  { width = 0.026667; value = 4012.713037; x = 0.180000; y = 0.340000 };
  { width = 0.026667; value = 4134.918332; x = 0.260000; y = 0.340000 };
  { width = 0.026667; value = 3705.954650; x = 0.340000; y = 0.340000 };
  { width = 0.026667; value = 3113.100990; x = 0.500000; y = 0.340000 };
  { width = 0.026667; value = 3581.639720; x = 0.580000; y = 0.340000 };
  { width = 0.026667; value = 3970.541725; x = 0.740000; y = 0.340000 };
```

```
  { width = 0.026667; value = 1994.007574; x = 0.820000; y = 0.340000 };
  { width = 0.026667; value = 2730.227002; x = 0.980000; y = 0.340000 };
  { width = 0.026667; value = 3046.871874; x = 0.060000; y = 0.420000 };
  { width = 0.026667; value = 4456.810333; x = 0.140000; y = 0.420000 };
  { width = 0.026667; value = 2851.158245; x = 0.220000; y = 0.420000 };
  { width = 0.026667; value = 2272.053455; x = 0.300000; y = 0.420000 };
  { width = 0.026667; value = 3461.120319; x = 0.380000; y = 0.420000 };
  { width = 0.026667; value = 3268.458853; x = 0.460000; y = 0.420000 };
  { width = 0.026667; value = 3082.460447; x = 0.540000; y = 0.420000 };
  { width = 0.026667; value = 3709.755859; x = 0.700000; y = 0.420000 };
  { width = 0.026667; value = 3161.854726; x = 0.020000; y = 0.500000 };
  { width = 0.026667; value = 3466.903882; x = 0.100000; y = 0.500000 };
  { width = 0.026667; value = 3716.447049; x = 0.180000; y = 0.500000 };
  { width = 0.026667; value = 3565.040993; x = 0.260000; y = 0.500000 };
  { width = 0.026667; value = 2918.063483; x = 0.340000; y = 0.500000 };
  { width = 0.026667; value = 2683.339196; x = 0.500000; y = 0.500000 };
  { width = 0.026667; value = 3331.351527; x = 0.660000; y = 0.500000 };
  { width = 0.026667; value = 2928.177514; x = 0.740000; y = 0.500000 };
  { width = 0.026667; value = 3021.576989; x = 0.900000; y = 0.500000 };
  { width = 0.026667; value = 3632.996958; x = 0.980000; y = 0.500000 };
  { width = 0.026667; value = 4317.129516; x = 0.060000; y = 0.580000 };
  { width = 0.026667; value = 3245.656047; x = 0.220000; y = 0.580000 };
  { width = 0.026667; value = 3275.257441; x = 0.300000; y = 0.580000 };
  { width = 0.026667; value = 3238.859563; x = 0.460000; y = 0.580000 };
  { width = 0.026667; value = 3540.446694; x = 0.540000; y = 0.580000 };
  { width = 0.026667; value = 3938.970629; x = 0.700000; y = 0.580000 };
  { width = 0.026667; value = 3582.033307; x = 0.860000; y = 0.580000 };
  { width = 0.026667; value = 5066.273236; x = 0.020000; y = 0.660000 };
  { width = 0.026667; value = 2570.051371; x = 0.260000; y = 0.660000 };
  { width = 0.026667; value = 2316.117427; x = 0.340000; y = 0.660000 };
  { width = 0.026667; value = 4523.811434; x = 0.420000; y = 0.660000 };
  { width = 0.026667; value = 3689.109567; x = 0.660000; y = 0.660000 };
  { width = 0.026667; value = 2372.421373; x = 0.740000; y = 0.660000 };
  { width = 0.026667; value = 3566.949827; x = 0.060000; y = 0.740000 };
  { width = 0.026667; value = 2696.655633; x = 0.220000; y = 0.740000 };
  { width = 0.026667; value = 3435.621790; x = 0.380000; y = 0.740000 };
  { width = 0.026667; value = 2376.047605; x = 0.460000; y = 0.740000 };
  { width = 0.026667; value = 3335.911108; x = 0.540000; y = 0.740000 };
  { width = 0.026667; value = 3452.314627; x = 0.620000; y = 0.740000 };
  { width = 0.026667; value = 2887.184716; x = 0.700000; y = 0.740000 };
  { width = 0.026667; value = 3610.375546; x = 0.780000; y = 0.740000 };
  { width = 0.026667; value = 2573.128731; x = 0.860000; y = 0.740000 };
  { width = 0.026667; value = 3415.345611; x = 0.020000; y = 0.820000 };
  { width = 0.026667; value = 3617.526764; x = 0.180000; y = 0.820000 };
  { width = 0.026667; value = 2789.440403; x = 0.340000; y = 0.820000 };
  { width = 0.026667; value = 3731.426877; x = 0.500000; y = 0.820000 };
  { width = 0.026667; value = 3713.099480; x = 0.580000; y = 0.820000 };
  { width = 0.026667; value = 3277.290639; x = 0.060000; y = 0.900000 };
  { width = 0.026667; value = 2987.352282; x = 0.300000; y = 0.900000 };
  { width = 0.026667; value = 4014.674202; x = 0.380000; y = 0.900000 };
  { width = 0.026667; value = 3888.595102; x = 0.460000; y = 0.900000 };
  { width = 0.026667; value = 3215.843482; x = 0.620000; y = 0.900000 };
  { width = 0.026667; value = 3248.446252; x = 0.700000; y = 0.900000 };
  { width = 0.026667; value = 3172.020630; x = 0.780000; y = 0.900000 };
  { width = 0.026667; value = 1786.763808; x = 0.860000; y = 0.900000 };
  { width = 0.026667; value = 4914.556562; x = 0.100000; y = 0.980000 };
  { width = 0.026667; value = 2941.888946; x = 0.260000; y = 0.980000 };
  { width = 0.026667; value = 2745.884646; x = 0.420000; y = 0.980000 };
  { width = 0.026667; value = 3384.683642; x = 0.900000; y = 0.980000 };
  { width = 0.026667; value = 4029.057164; x = 0.980000; y = 0.980000 };
  ] } }

| init psB {spatial = { points = [
  { width = 0.026667; value = 1; x = 0.100000; y = 0.020000 };
  { width = 0.026667; value = 1; x = 0.180000; y = 0.020000 };
  { width = 0.026667; value = 1; x = 0.340000; y = 0.020000 };
  { width = 0.026667; value = 1; x = 0.420000; y = 0.020000 };
  { width = 0.026667; value = 1; x = 0.500000; y = 0.020000 };
  { width = 0.026667; value = 1; x = 0.740000; y = 0.020000 };
  { width = 0.026667; value = 1; x = 0.900000; y = 0.020000 };
  { width = 0.026667; value = 1; x = 0.060000; y = 0.100000 };
  { width = 0.026667; value = 1; x = 0.140000; y = 0.100000 };
  { width = 0.026667; value = 1; x = 0.220000; y = 0.100000 };
  { width = 0.026667; value = 1; x = 0.380000; y = 0.100000 };
  { width = 0.026667; value = 1; x = 0.460000; y = 0.100000 };
  { width = 0.026667; value = 1; x = 0.780000; y = 0.100000 };
```

```
  { width = 0.026667; value = 1; x = 0.860000; y = 0.100000 };
  { width = 0.026667; value = 1; x = 0.100000; y = 0.180000 };
  { width = 0.026667; value = 1; x = 0.500000; y = 0.180000 };
  { width = 0.026667; value = 1; x = 0.580000; y = 0.180000 };
  { width = 0.026667; value = 1; x = 0.660000; y = 0.180000 };
  { width = 0.026667; value = 1; x = 0.740000; y = 0.180000 };
  { width = 0.026667; value = 1; x = 0.900000; y = 0.180000 };
  { width = 0.026667; value = 1; x = 0.300000; y = 0.260000 };
  { width = 0.026667; value = 1; x = 0.380000; y = 0.260000 };
  { width = 0.026667; value = 1; x = 0.540000; y = 0.260000 };
  { width = 0.026667; value = 1; x = 0.620000; y = 0.260000 };
  { width = 0.026667; value = 1; x = 0.780000; y = 0.260000 };
  { width = 0.026667; value = 1; x = 0.860000; y = 0.260000 };
  { width = 0.026667; value = 1; x = 0.940000; y = 0.260000 };
  { width = 0.026667; value = 1; x = 0.100000; y = 0.340000 };
  { width = 0.026667; value = 1; x = 0.420000; y = 0.340000 };
  { width = 0.026667; value = 1; x = 0.660000; y = 0.340000 };
  { width = 0.026667; value = 1; x = 0.900000; y = 0.340000 };
  { width = 0.026667; value = 1; x = 0.620000; y = 0.420000 };
  { width = 0.026667; value = 1; x = 0.780000; y = 0.420000 };
  { width = 0.026667; value = 1; x = 0.860000; y = 0.420000 };
  { width = 0.026667; value = 1; x = 0.940000; y = 0.420000 };
  { width = 0.026667; value = 1; x = 0.420000; y = 0.500000 };
  { width = 0.026667; value = 1; x = 0.580000; y = 0.500000 };
  { width = 0.026667; value = 1; x = 0.820000; y = 0.500000 };
  { width = 0.026667; value = 1; x = 0.140000; y = 0.580000 };
  { width = 0.026667; value = 1; x = 0.380000; y = 0.580000 };
  { width = 0.026667; value = 1; x = 0.620000; y = 0.580000 };
  { width = 0.026667; value = 1; x = 0.780000; y = 0.580000 };
  { width = 0.026667; value = 1; x = 0.940000; y = 0.580000 };
  { width = 0.026667; value = 1; x = 0.100000; y = 0.660000 };
  { width = 0.026667; value = 1; x = 0.180000; y = 0.660000 };
  { width = 0.026667; value = 1; x = 0.500000; y = 0.660000 };
  { width = 0.026667; value = 1; x = 0.580000; y = 0.660000 };
  { width = 0.026667; value = 1; x = 0.820000; y = 0.660000 };
  { width = 0.026667; value = 1; x = 0.900000; y = 0.660000 };
  { width = 0.026667; value = 1; x = 0.980000; y = 0.660000 };
  { width = 0.026667; value = 1; x = 0.140000; y = 0.740000 };
  { width = 0.026667; value = 1; x = 0.300000; y = 0.740000 };
  { width = 0.026667; value = 1; x = 0.940000; y = 0.740000 };
  { width = 0.026667; value = 1; x = 0.100000; y = 0.820000 };
  { width = 0.026667; value = 1; x = 0.260000; y = 0.820000 };
  { width = 0.026667; value = 1; x = 0.420000; y = 0.820000 };
  { width = 0.026667; value = 1; x = 0.660000; y = 0.820000 };
  { width = 0.026667; value = 1; x = 0.740000; y = 0.820000 };
  { width = 0.026667; value = 1; x = 0.820000; y = 0.820000 };
  { width = 0.026667; value = 1; x = 0.900000; y = 0.820000 };
  { width = 0.026667; value = 1; x = 0.980000; y = 0.820000 };
  { width = 0.026667; value = 1; x = 0.140000; y = 0.900000 };
  { width = 0.026667; value = 1; x = 0.220000; y = 0.900000 };
  { width = 0.026667; value = 1; x = 0.540000; y = 0.900000 };
  { width = 0.026667; value = 1; x = 0.940000; y = 0.900000 };
  { width = 0.026667; value = 1; x = 0.020000; y = 0.980000 };
  { width = 0.026667; value = 1; x = 0.180000; y = 0.980000 };
  { width = 0.026667; value = 1; x = 0.340000; y = 0.980000 };
  { width = 0.026667; value = 1; x = 0.500000; y = 0.980000 };
  { width = 0.026667; value = 1; x = 0.580000; y = 0.980000 };
  { width = 0.026667; value = 1; x = 0.660000; y = 0.980000 };
  { width = 0.026667; value = 1; x = 0.740000; y = 0.980000 };
  { width = 0.026667; value = 1; x = 0.820000; y = 0.980000 };
  ] } }

| init F2Q2 {spatial = { points = [
  { width = 0.026667; value = 5804.447107; x = 0.100000; y = 0.020000 };
  { width = 0.026667; value = 6122.649409; x = 0.180000; y = 0.020000 };
  { width = 0.026667; value = 4891.526801; x = 0.340000; y = 0.020000 };
  { width = 0.026667; value = 5649.905421; x = 0.420000; y = 0.020000 };
  { width = 0.026667; value = 3782.217583; x = 0.500000; y = 0.020000 };
  { width = 0.026667; value = 4499.596078; x = 0.740000; y = 0.020000 };
  { width = 0.026667; value = 5383.485388; x = 0.900000; y = 0.020000 };
  { width = 0.026667; value = 4370.039519; x = 0.060000; y = 0.100000 };
  { width = 0.026667; value = 3329.871407; x = 0.140000; y = 0.100000 };
  { width = 0.026667; value = 4098.420337; x = 0.220000; y = 0.100000 };
  { width = 0.026667; value = 5455.747875; x = 0.380000; y = 0.100000 };
  { width = 0.026667; value = 5610.996546; x = 0.460000; y = 0.100000 };
  { width = 0.026667; value = 4901.707403; x = 0.780000; y = 0.100000 };
```

```
  { width = 0.026667; value = 4110.247127; x = 0.860000; y = 0.100000 };
  { width = 0.026667; value = 4050.574502; x = 0.100000; y = 0.180000 };
  { width = 0.026667; value = 2962.133629; x = 0.500000; y = 0.180000 };
  { width = 0.026667; value = 3454.625662; x = 0.580000; y = 0.180000 };
  { width = 0.026667; value = 6195.222333; x = 0.660000; y = 0.180000 };
  { width = 0.026667; value = 5333.679290; x = 0.740000; y = 0.180000 };
  { width = 0.026667; value = 5033.114851; x = 0.900000; y = 0.180000 };
  { width = 0.026667; value = 4978.723396; x = 0.300000; y = 0.260000 };
  { width = 0.026667; value = 5205.275878; x = 0.380000; y = 0.260000 };
  { width = 0.026667; value = 4200.936241; x = 0.540000; y = 0.260000 };
  { width = 0.026667; value = 4649.418839; x = 0.620000; y = 0.260000 };
  { width = 0.026667; value = 4792.228072; x = 0.780000; y = 0.260000 };
  { width = 0.026667; value = 4589.119578; x = 0.860000; y = 0.260000 };
  { width = 0.026667; value = 4083.372369; x = 0.940000; y = 0.260000 };
  { width = 0.026667; value = 5043.202876; x = 0.100000; y = 0.340000 };
  { width = 0.026667; value = 4725.786435; x = 0.420000; y = 0.340000 };
  { width = 0.026667; value = 5920.534994; x = 0.660000; y = 0.340000 };
  { width = 0.026667; value = 4977.463544; x = 0.900000; y = 0.340000 };
  { width = 0.026667; value = 5864.960746; x = 0.620000; y = 0.420000 };
  { width = 0.026667; value = 5382.316436; x = 0.780000; y = 0.420000 };
  { width = 0.026667; value = 4756.539756; x = 0.860000; y = 0.420000 };
  { width = 0.026667; value = 5357.569437; x = 0.940000; y = 0.420000 };
  { width = 0.026667; value = 4507.272383; x = 0.420000; y = 0.500000 };
  { width = 0.026667; value = 5977.897025; x = 0.580000; y = 0.500000 };
  { width = 0.026667; value = 5671.619173; x = 0.820000; y = 0.500000 };
  { width = 0.026667; value = 5920.839392; x = 0.140000; y = 0.580000 };
  { width = 0.026667; value = 6878.735430; x = 0.380000; y = 0.580000 };
  { width = 0.026667; value = 5334.460155; x = 0.620000; y = 0.580000 };
  { width = 0.026667; value = 5228.356644; x = 0.780000; y = 0.580000 };
  { width = 0.026667; value = 5330.781644; x = 0.940000; y = 0.580000 };
  { width = 0.026667; value = 5861.712008; x = 0.100000; y = 0.660000 };
  { width = 0.026667; value = 4181.884322; x = 0.180000; y = 0.660000 };
  { width = 0.026667; value = 4237.059662; x = 0.500000; y = 0.660000 };
  { width = 0.026667; value = 5972.961948; x = 0.580000; y = 0.660000 };
  { width = 0.026667; value = 6960.910529; x = 0.820000; y = 0.660000 };
  { width = 0.026667; value = 5551.375668; x = 0.900000; y = 0.660000 };
  { width = 0.026667; value = 6766.606498; x = 0.980000; y = 0.660000 };
  { width = 0.026667; value = 5194.109685; x = 0.140000; y = 0.740000 };
  { width = 0.026667; value = 5058.039087; x = 0.300000; y = 0.740000 };
  { width = 0.026667; value = 4472.065713; x = 0.940000; y = 0.740000 };
  { width = 0.026667; value = 4442.015433; x = 0.100000; y = 0.820000 };
  { width = 0.026667; value = 4674.247628; x = 0.260000; y = 0.820000 };
  { width = 0.026667; value = 5914.932745; x = 0.420000; y = 0.820000 };
  { width = 0.026667; value = 5678.678119; x = 0.660000; y = 0.820000 };
  { width = 0.026667; value = 5989.529588; x = 0.740000; y = 0.820000 };
  { width = 0.026667; value = 5971.749582; x = 0.820000; y = 0.820000 };
  { width = 0.026667; value = 5054.292376; x = 0.900000; y = 0.820000 };
  { width = 0.026667; value = 4590.416176; x = 0.980000; y = 0.820000 };
  { width = 0.026667; value = 6208.487599; x = 0.140000; y = 0.900000 };
  { width = 0.026667; value = 3873.851092; x = 0.220000; y = 0.900000 };
  { width = 0.026667; value = 4547.028556; x = 0.540000; y = 0.900000 };
  { width = 0.026667; value = 4612.411055; x = 0.940000; y = 0.900000 };
  { width = 0.026667; value = 5318.234381; x = 0.020000; y = 0.980000 };
  { width = 0.026667; value = 3620.071058; x = 0.180000; y = 0.980000 };
  { width = 0.026667; value = 4652.524274; x = 0.340000; y = 0.980000 };
  { width = 0.026667; value = 4376.188481; x = 0.500000; y = 0.980000 };
  { width = 0.026667; value = 3177.612205; x = 0.580000; y = 0.980000 };
  { width = 0.026667; value = 4914.247698; x = 0.660000; y = 0.980000 };
  { width = 0.026667; value = 4525.008686; x = 0.740000; y = 0.980000 };
  { width = 0.026667; value = 5534.780730; x = 0.820000; y = 0.980000 };
  ] } }

| init ActOut 100.000000
| init FuelOut 1000.000000

| psA + ActOut <->{(3/rprot)*P}{(3/rprot)*P} ActA + psA
| ActA + F1Q1 ->{kdis} F1Act + Q1A
| psA + Q1A <->{(3/rprot)*P}{(3/rprot)*P} Q1sol + psA

| psB + Q1sol <->{(3/rprot)*P}{(3/rprot)*P} Q1B + psB
| psB + FuelOut <->{(3/rprot)*P}{(3/rprot)*P} FuelB + psB
| Q1B + F2Q2 ->{kdis} F2Q1 + Q2B
| F2Q1 + FuelB ->{kFuel} F2Fuel + Q1B
| psB + Q2B <->{(3/rprot)*P}{(3/rprot)*P} Q2sol + psB
```

#### Supplementary Tables

##### Supplementary table 1 | DNA sequences of the simple compartmentalized DSD reaction.

| | | Length # bases | 5' modification | 3' modification |
|---|---|---|---|---|
| **Fig. 1d** | | | | |
| F ($F_1$) | GTAGTAGT GCATTAGT GTCG TTCGTTCG CTGTAATA | 36 | Cy5 | Biotin-TEG |
| Q ($Q_1$) | CGAACGAA CGAC CATGATAG ACTAATGC ACTACTAC | 36 | | Iowa Black |
| A (Input) | TATTACAG CGAACGAA CGAC ACTAATGC ACTACTAC | 36 | | |
| **Fig. S5c** | | | | |
| A_Alexa546 | TATTACAG CGAACGAA CGAC ACTAATGC ACTACTAC | 36 | Alexa546 | |
| A_NonComp_Cy3 | AGACCGGT CGAACGAA CGAC ACTAATGC ACTACTAC | 36 | Cy3 | |

##### Supplementary table 2 | DNA sequences of two-layer signaling cascade.

| | | Length # bases | 5' modification | 3' modification |
|---|---|---|---|---|
| **Fig. Fig. 2a, and Fig. S7a** | | | | |
| F ($F_1$) | GTAGTAGT GCATTAGT GTCG TTCGTTCG CTGTAATA | 36 | Cy5 | Biotin-TEG |
| Q ($Q_1$) | CGAACGAA CGAC CATGATAG ACTAATGC ACTACTAC | 36 | | Iowa Black |
| A (Input) | TATTACAG CGAACGAA CGAC ACTAATGC ACTACTAC | 36 | | |
| $F_2$ | GCATTAGT CTATCATG GTCG TTCGTTCG ACAGTTCC | 36 | Biotin-TEG | Alexa546 |
| $Q_2$ | GGAACTGT CGAACGAA CGTGAAAC CGAC CATGATAG | 36 | Iowa Black | phosphate |
| Fuel | GGAACTGT CGAACGAA CGAC CATGATAG | 28 | | phosphate |
| **Fig. S7b** | | | | |
| $Q_1$_Cy3 | CGAACGAA CGAC CATGATAG ACTAATGC ACTACTAC | 36 | Cy3 | Iowa Black |

##### Supplementary table 3 | DNA sequences of three-layer signaling cascade.

| | | Length # bases | 5' modification | 3' modification |
|---|---|---|---|---|
| **Fig. 2g and Fig. S8a** | | | | |
| $F_1$ | GTGTATG AGTATGG TTGGTGA AAAGGTA GTGAGAT TTTTT | 40 | Cy5 | Biotin-TEG |
| $Q_1$ | TACCTTT TCACCAA TCACCTT CCATACT CATACAC | 35 | | Iowa Black |
| $F_2$ | TTTTT AGTATGG AAGGTGA TTGGTGA AAAGGTA GGTTTTG | 40 | Biotin-TEG | Cy5 |
| $Q_2$ | CAAAACC TACCTTT ACCTCTA TCACCAA TCACCTT | 35 | Iowa Black | |
| $F_3$ | GAATGTG AAGGTGA TTGGTGA TAGAGGT AAAGGTA | 35 | Cy5 | Biotin-TEG |

| | | | | |
|---|---|---|---|---|
| $Q_3$ | ACCTCTA TCACCAA CTTATCC TCACCTT CACATTC | 35 | | Iowa Black |
| Input | ATCTCAC TACCTTT TCACCAA CCATACT | 28 | | |
| Fuel1 | TACCTTT TCACCAA CCATACT CATACAC | 28 | | |
| Fuel2 | CAAAACC TACCTTT TCACCAA TCACCTT | 28 | | |
| Fuel3 | ACCTCTA TCACCAA TCACCTT CACATTC | 28 | | |

**Supplementary table 4 | DNA sequences of the negative feedback system.**

| | | Length # bases | 5' modification | 3' modification |
|---|---|---|---|---|
| **Fig. 3a and Fig. S9a** | | | | |
| F1 | GTAGTA GTGGATTA GTGTAG TTAGTTAG GTGTAATA | 36 | Cy5 | Biotin-TEG |
| Q1 | TTCA CTAACTAA CTACAC ATACTTAC TAATCCAC TACTAC | 40 | | Iowa Black |
| F2 | TTTTTTTT TAAGTAT GTGTAG TTAGTTAG TGAA AGAGTTG | 40 | Biotin-TEG | Cy3 |
| Inh | CAACTCT TTCA TATTACAC CTAACTAA CTACAC TAATCCAC TACTAC | 47 | Iowa Black | Iowa Black |
| Input | TATTACAC CTAACTAA CTACAC TAATCCAC | 30 | | |
| Fuel | AACTCT TTCA CTAACTAA CTACAC | 24 | | |

**Supplementary table 5 | DNA sequences of the logic circuits.**

| | | Length # bases | 5' modification | 3' modification |
|---|---|---|---|---|
| **Fig. 3d AND gate and Fig. S10a** | | | | |
| F1 | AGTTGGATGAGAGGGATGTGGTGA | 24 | Cy5 | Biotin-TEG |
| Signal A | TCCCTCTCATCCAACTTCTCTCTACTATTTCATTTTCATT | 40 | | |
| Input A | TCACCACATCCCTCTCATCCAACT | 24 | | Iowa Black |
| F2 | AGGTAGGTGAGATGAGTGTATAGG | 24 | Cy5 | Biotin-TEG |
| Signal B | CTCATCTCACCTACCTACTAACTCAATACTAAAACTAACA | 40 | | |
| Input B | CCTATACACTCATCTCACCTACCT | 24 | | Iowa Black |
| F3 | TGTTAGTTTTAGTATTGAGTTAGTAATGAAAATGAAATAGTAGAGAGA | 48 | Cy5 | Biotin-TEG |
| Blocker | CTATTTCATTTTCATTACTAACTC | 24 | | |
| Output | AATACTAAAACTAACA | 16 | | Iowa Black |
| **Fig. 3d OR gate and Fig. S10b** | | | | |
| F1 | AGGTAGGTTGAGGTGTGAAGGATT | 24 | Cy5 | Biotin-TEG |
| Signal A | ACACCTCAACCTACCTTTTCACTCCTCCTAAA | 40 | | |
| Input A | AATCCTTCACACCTCA ACCTACCT | 24 | | Iowa Black |
| F2 | AGGTAGGTGAGATGAGTGTATAGG | 24 | Cy5 | Biotin-TEG |
| Signal B | CTCATCTCACCTACCTTTTCACTCCTCCTAAA | 40 | | |
| Input B | CCTATACACTCATCTC ACCTACCT | 24 | | Iowa Black |
| F3 | TTTAGGAGGAGTGAAAAGGTAGGT | 24 | Cy5 | Biotin-TEG |
| Output | TTTCACTCCTCCTAAA | 16 | | Iowa Black |
| **Fig. 3f** | | | | |
| B OR C | TTTCACTCCTCCTAAA ACTAACTC AATACTAAAACTAACA | 40 | | |
| Input B | ACACCTCACCCACTCT ACCTACCT TTTCACTCCTCCTAAA | 40 | | |
| Input C | CTCATCTCATACTATC ACCTACCT TTTCACTCCTCCTAAA | 40 | | |

**Supplementary table 6 | DNA sequences of experiments in FBS.**

| | | Length # bases | 5' modification | 3' modification |
|---|---|---|---|---|
| **Fig. 4a, Fig. 4c and Fig. 4e** | | | | |

| | | | | |
|---|---|---|---|---|
| F ($F_1$) | GTAGTAGTGCATTAGTGTCGTTCGTTCGCTGTAATA | 36 | Cy5 | Biotin-TEG |
| Q (Signal $Q_1$) | CGAACGAACGACCATGATAGACTAATGCACTACTAC | 36 | | Iowa Black |
| A (Input) | TATTACAGCGAACGAACGACACTAATGCACTACTAC | 36 | | |
| $F_2$ | GTAGTAGTGCATTAGTCTATCATGGTCGTTCGTTCG | 36 | Biotin-TEG | Alexa546 |
| $Q_2$ | CGAACGAACGACCATGCGTGAAACATAGACTAATGC | 36 | Iowa Black | phosphate |
